# Prototypical pacemaker neurons are immunocompetent cells

**DOI:** 10.1101/750026

**Authors:** Alexander Klimovich, Stefania Giacomello, Åsa Björklund, Louis Faure, Marketa Kaucka, Christoph Giez, Andrea P. Murillo-Rincon, Ann-Sophie Matt, Gabriele Crupi, Jaime de Anda, Gerard C.L. Wong, Mauro D’Amato, Igor Adameyko, Thomas C.G. Bosch

## Abstract

Pacemaker neurons exert control over neuronal circuit function by their intrinsic ability to generate rhythmic bursts of action potential. Recent work has identified rhythmic gut contractions in human, mice and hydra to be dependent on both neurons and the resident microbiota. However, little is known about the evolutionary origin of these neurons and their interaction with microbes. In this study, we identified and functionally characterized prototypical ANO/SCN/TRPM ion channel expressing pacemaker cells in the basal metazoan *Hydra* by using a combination of single-cell transcriptomics, immunochemistry, and functional experiments. Unexpectedly, these prototypical pacemaker neurons express a rich set of immune-related genes mediating their interaction with the microbial environment. Functional experiments validated a model of the evolutionary emergence of pacemaker cells as neurons using components of innate immunity to interact with the microbial environment and ion channels to generate rhythmic contractions.

---

The enteric nervous system (ENS) coordinates the major functions of the gastrointestinal tract^1^. In all extant animals, the structurally conserved ENS is a diffuse nerve net located within the wall of the gastrointestinal tract. In pre-bilaterian animals, such as *Hydra* the nervous system is structurally simple and thus has great potential to inform us about evolutionary ancient basic structural and functional principles of neural circuits^2^ (Fig. 1a). The principal function of the ENS is coordination of the rhythmic intestine motility, known as peristalsis, that occurs ubiquitously in the animal kingdom (Fig. 1a) and is driven by rhythmic electrical pulses generated by pacemaker cells^3^. In mammals, proper functioning of the interstitial cells of Cajal (ICC), that serve as pacemakers in the muscular wall of the gastrointestinal tract^4–6^, is essential for normal gut motility^7–9^. Dysfunction of the pacemaker system contributes to functional gastrointestinal disorders (FGID) such as irritable bowel syndrome (IBS), chronic constipation and intestinal pseudo-obstruction^9–12^. The Ca^2+^-activated Cl^−^-channel Anoctamin-1 (encoded by *ANO1* gene) and the voltage-gated Na^+^-channel Nav1.5 (*SCN5A* gene) represent molecular markers of the interstitial pacemaker cells in human and mice^13–15^ and DNA variants in the corresponding genes have been shown to associate with increased risk of IBS^16–18^. Ion channel dysfunction (channelopathy) appears to be a plausible pathogenetic mechanism in FGIDs^19^, as the transient receptor potential cation channel TRPM8 (known as the cold and menthol receptor) and other ion channels have also been implicated in IBS susceptibility and gut dysmotility^20–23^. Spontaneous contractile gut activities not only are affected by microbes. In fact, there is evidence that bacterial population dynamics themselves are affected by the periodic stimulation^24^. Previous studies in *Hydra* suggested that the rhythmic peristaltic movements of the body column are dependent on neurons^25^ and that they are modulated by the host-associated microbiota since germ-free animals display reduced and less regular contraction frequencies^26^ Single-cell RNA sequencing uncovered neuron specific transcriptional signatures and the presence of distinct neuronal subtypes^27^. Little, however, little is known about the nature of the neurons that generate peristaltic movement in a pre-bilaterian animal and how such prototypical neurons engage with the resident microbiota.

**Fig. 1.**
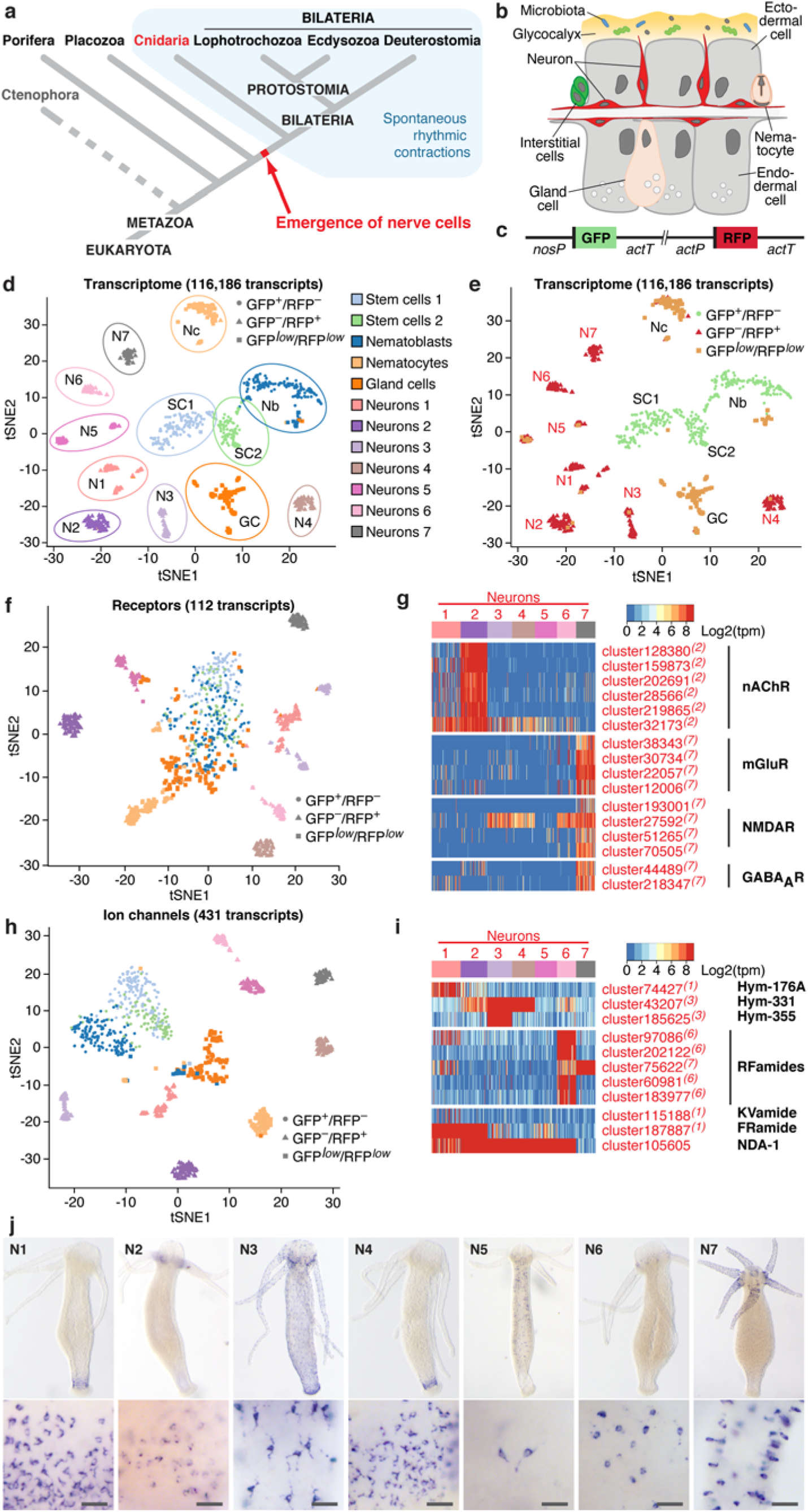
Single-cell transcriptome profiling uncovers the molecular anatomy of *Hydra* nervous system. **a**, Emergence of the first nerve cells preceded the divergence of Cnidaria and Bilateria. Cnidarians possess structurally simple nervous systems and offer a great potential to reveal the fundamental structural and functional principles of neural circuits. Spontaneous rhythmic contractions are ubiquitously observed in Eumetazoa. **b**, The *Hydra* body is made of three cell lineages: the ectodermal and endodermal epithelia separated by the extracellular matrix, and the lineage of interstitial cells. The outer surface of the ectoderm is covered by a glycocalyx that serves as a habitat for symbiotic bacteria. The endoderm lining the gastric cavity, is free of glycocalyx and stable microbiota. Two nerve nets made of sensory and ganglion neurons are embedded within the both epithelia. **c**, Genetic construct used to generate transgenic *Hydra* polyps and differentially label cells within the interstitial lineage by a combination of two fluorescent proteins. Green fluorescent protein (GFP) was expressed under a stem-cell specific *nanos* promoter (nos-P), the red fluorescent protein (RFP) was driven by the *actin* promoter (act-P) that is particularly active in terminally differentiated neurons. Both cassettes are flanked by the *actin* terminator (act-T). **d**, t-SNE map constructed by dimensionality-reduction principal component analysis defined by highly covariable genes (see Methods). Total 928 cells were partitioned in 12 clusters and colored by their cell-type identities inferred from expressed proliferation- and cell type-specific marker genes (see Supplementary Data 5 and 6). **e**, t-SNE map based on analysis of the entire reference transcriptome made of 116,186 transcripts clearly segregates 12 clusters, including 7 subpopulations of neurons. Cells are color-coded by their phenotype captured by FACS upon sorting. **f**, t-SNE map based on expression analysis of 112 transcripts coding for putative neurotransmitter receptors (see Supplementary Data 7). While stem cells and nematoblasts, nematocytes and gland cells form one cluster, neuronal populations are clearly segregated, indicating that each neuronal population is characterized by a specific set of receptors. **g**, Heatmap illustrating expression of genes coding for putative nicotinic acetylcholine receptors (nAChR), muscarinic and N-methyl-D-aspartate glutamate receptors (mGluR and NMDAR), and γ-aminobutyric acid type-A receptors (GABAAR) within seven neuronal populations. Expression within the entire interstitial lineage is presented in Extended Data Fig. 3. Transcripts specifically upregulated in the neurons are labelled red, superscript numbers indicate the nerve cell cluster (N1-N7) where the transcripts are significantly (padj<0.05) enriched. **h**, t-SNE map constructed by expression analysis of 431 transcripts coding for putative ion channels (see Supplementary Data 8). Seven neuronal populations are clearly segregated, suggesting that each neuronal population is characterized by a specific set of channels. **i**, Heatmap illustrates expression of 11 genes coding for main known neuropeptides in *Hydra*. Each neuronal population expresses a unique combination of neuropeptides. **j**, *In situ* hybridization for marker genes strongly enriched in each of seven nerve cells clusters (N1-N7, see Extended Data Fig. 6) reveals that seven neuronal subpopulations reside in spatially restricted domains along the body column of *Hydra*. Scale bar: 25 µm.

Here, we provide a definition of prototypical pacemaker cells, which integrates marker genes discovered in human dysmotility patients with the recent discovery that spontaneous contractile activities are affected by microbes. The functional experiments connected the rhythm generation and interactions with microbes at the level of this specific neuronal population. These findings shed new light on the evolution of pacemaker neurons, emphasize the role of the microbial environment in dysmotility, and underscore the importance of cross-species comparisons in tracking cell type evolution.

## Identification of pacemaker cells in *Hydra* using human orthologous genes

Previous studies in *Hydra* suggested that a subpopulation of neurons located in the head region^28–30^ might have properties of pacemaker cells to control the regularity of spontaneous body contractions. To gain insight into this specific cell population, we first assessed the molecular and functional diversity of the neuronal populations by single cell RNA-sequencing using RFP-labelled neurons (Fig. 1b–j, Extended Data Fig. 1 and 2). Similar to the analysis of the entire transcriptome, the expression profile of 112 transcripts coding for putative neurotransmitter receptors clearly identified seven distinct clusters of neurons and separated them from the stem cells and non-neuronal cells (Fig. 1d–f). This indicates that each neuronal subpopulation is characterized by a specific set of neurotransmitter receptors (Fig. 1g, Extended Data Fig. 3). For instance, most transcripts coding for putative nicotinic acetylcholine receptors (nAChRs) were expressed almost exclusively in neuronal subpopulation N2 (Fig. 1g, Extended Data Fig. 3), while diverse homologues of muscarinic and N-methyl-D-aspartate glutamate receptors (mGluRs and NMDARs), and γ-aminobutyric acid type-A receptors (GABAARs) were enriched in the neuronal subpopulation N7 (Fig. 1g, Extended Data Fig. 3). Similarly, the expression profiles of 431 transcripts coding for ion channels clearly segregated GFP^+^/RFP^−^ stem cells from the GFP*^low^*/RFP*^low^* non-neuronal cells and the seven remarkably distinct clusters of GFP^−^/RFP^+^ neurons (Fig. 1h). In addition, each neuronal subpopulation was found to express a unique combination of neuropeptides (Fig. 1i, Extended Data Fig. 4). The Hym-355 neuropeptide precursor gene, for instance, was exclusively expressed in the neuronal subpopulation N3, while Hym-176A transcripts were discovered predominantly in the N1 subpopulation. Most of RFamide precursor transcripts were found in subpopulation N6 (Fig. 1i, Extended Data Fig. 4). Genes coding for some other peptides, including Hym-331 and FRamide, were expressed in two or more neuronal subpopulations. Taken together, these observations indicate that the molecular identity of different subpopulations of neurons is determined by the specific expression of ion channels, neurotransmitter receptors and neuropeptides.

Interestingly, based on the expression profiles of 364 transcripts coding for putative transcription factors (TFs), all seven subpopulations of neurons express a common set of TFs, which separates them from stem cells and non-neuronal cell types (Extended Data Fig. 5). The neuron-specific TF signature consists mainly of Zn-finger, homeodomain and helix-loop-helix DNA-binding proteins, including the Achaete-scute homologous TFs. Further, each neuronal population is characterized by a combinatorial expression of few genes encoding other TFs, such as the homologues of Aristales, NeuroD, and Orthopedia (Extended Data Fig. 5b, c, Supplementary Data 1).

*In situ* hybridization for selected marker genes enriched in each of the seven neuronal subpopulations (Fig. 1j, Extended Data Fig. 6a) showed that the neuronal subpopulations in *Hydra* reside in spatially restricted domains along the body column. Neuronal clusters N1 were found confined to the foot region of the polyp (Fig. 1j, Extended Data Fig. 6b). Cluster N2 was represented by a population of neurons located in the base of tentacles. N3-specific neurons were spread in the ectoderm along the entire *Hydra* body, whereas neurons of subpopulations N4 and N5 were found in the endodermal epithelial layer (Fig. 1j, Extended Data Fig. 6b). Neurons from the cluster N6 were confined to the hypostome of a polyp, whereas neurons of the cluster N7 were restricted to the tentacles. Notably, most of the seven spatially restricted neuronal subpopulations contain both sensory and ganglion cells (Fig. 1j). Recently, a similar molecular map of the *Hydra* nervous system was published^30^. The higher number of cells sequenced (∼20,000) allowed Siebert and co-authors to perform deeper clustering of the neuronal population and uncover 12 differentiated neuronal subtypes. Analysis of marker gene expression (Extended Data Fig. 7) indicated a clear correspondence between the neuronal clusters identified buy us and the subtypes reported by Siebert and co-authors^27^.

To identify the pacemaker cells among the neurons in *Hydra*, we focused on a few human orthologs known to be either restricted in their expression to human ICCs (ANO1 and SCN5A)^13,14^ or mechanistically involved in the control of gut motility via circular smooth muscle cell contractions (menthol sensitive Ca2+ channels such as TRPM8)^31^ (Fig. 2a). Homology search and phylogenetic analysis uncovered three *Hydra* genes coding for SCN-like sodium channels, six homologues of ANO1-like chloride channels and four homologues encoding TRPM-like cation channels, which are remarkably similar to their human counterparts (Fig. 2b–d, Extended Data Fig. 8–10). Analysis of the single cell transcriptome (Fig. 2e, Extended Data Fig. 11) revealed that the expression of genes encoding SCN-, ANO1- and TRPM-like channels overall was very weak, often restricted to only few cells. However, several of the transcripts coding for SCN- and ANO1-homologues were more expressed in neurons (Fig. 2e, Extended Data Fig. 11) with some of the transcripts specific for neuronal subpopulation N2, which is located at the base of tentacles (Fig. 1j and 2e, Extended Data Fig. 11). Real-time PCR confirmed (Fig. 2f) that most of the SCN-, ANO1- and TRPM-like ion channel genes are upregulated in the head region. Transcripts of two of these genes, *cluster2505* and *cluster30856*, coding for ANO1-like and SCN-like channels, could be detected by *in situ* hybridization at the base of tentacles (Fig. 2g, h). Immunohistochemical analysis using specific antibodies raised against synthetic peptides confirmed the presence of ANO1-like and SCN-like channel proteins at the base of the tentacles (Fig. 2i–k) in the subpopulation N2 domain (Fig. 1j). High magnification confocal microscopy identified the cells expressing SCN- and ANO1- channels as neurons (Fig. 2l, m). Taken together, these observations indicate that the genes encoding SCN- and ANO1- and TRPM-like channels are expressed in a population of nerve cells resident in the subpopulation N2 at the base of tentacles (Fig. 2n).

**Fig. 2.**
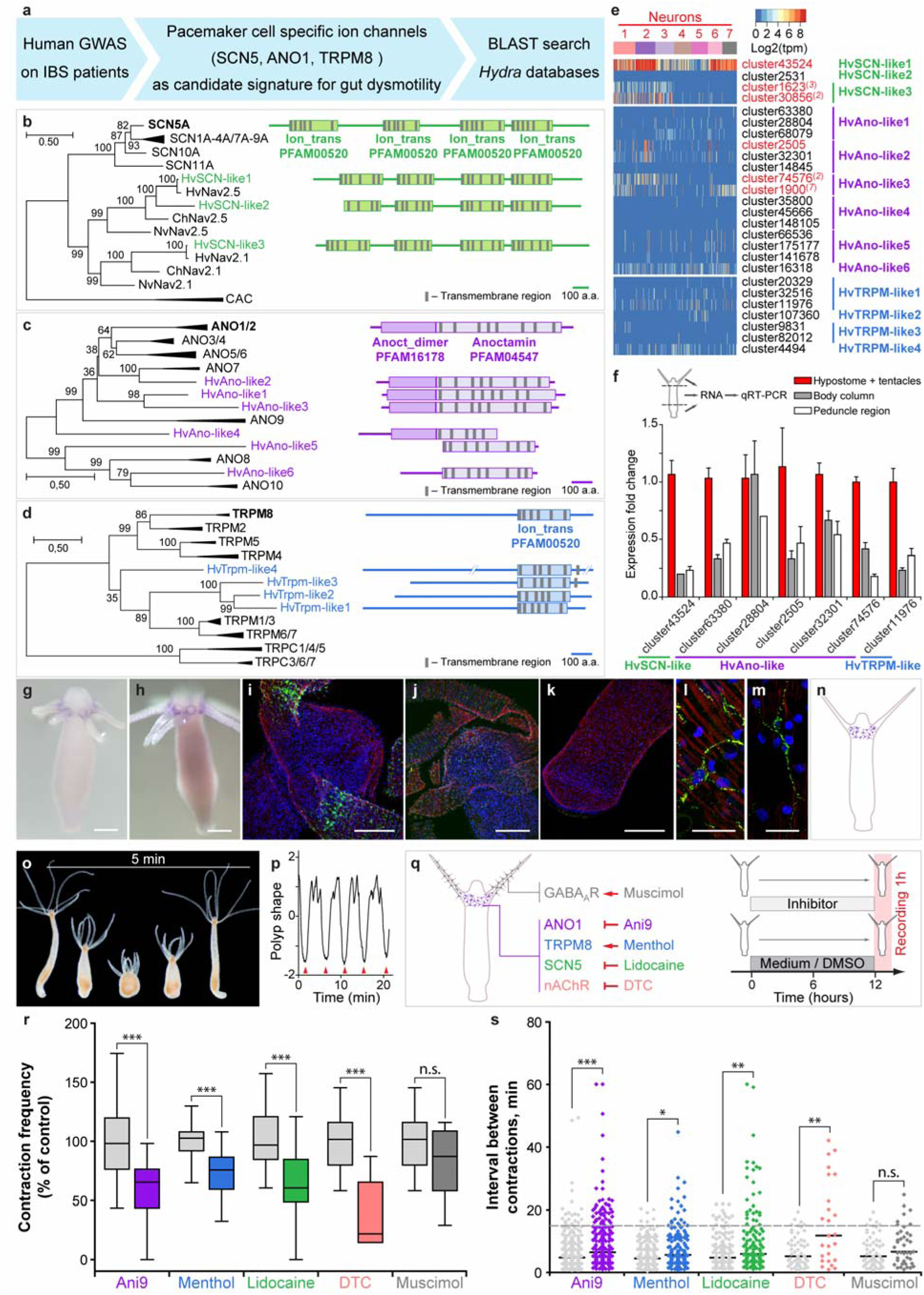
Identification of *Hydra* pacemaker cells using orthologs of human ion channels. **a**, Genome-wide association studies on patients with gut motility disorders such as irritable bowel syndrome (IBS) identified a set of ion channels SCN5, ANO1 and TRPM8 that are expressed in human pacemaker cells (ICCs) and found to be essential for gut motility control. BLAST search was used to identify the homologous genes in *Hydra*. **b–d**, Pacemaker-specific ion channels are highly conserved in *Hydra*. Phylogenetic tree and domain structure of human SCN (on B), ANO1 (on c) and TRPM (on d) channels and the orthologs from *Hydra* (Hv). Additionally, sequences from other cnidarians, *Nematostella vectensis* (Nv) and *Clytia hemisphaerica* (Ch) are included to the phylogenetic analysis. Non-collapsed trees are presented on Extended Data Fig. 8–10. The topology and domain structure of three *Hydra* SCN-like sodium channels, six ANO1-like chloride channels and four homologues of TRPM-like cation channels are remarkably similar to their human counterparts. **e**, Expression of genes encoding SCN-, ANO1- and TRPM-like channels in *Hydra* single-cell dataset is overall very weak, and restricted to only few cells. However, several transcripts coding for SCN- and ANO1-homologues are significantly up-regulated in neurons (red) with some of the transcripts specifically enriched in neuronal subpopulation N2 (red superscript). **f**, Expression levels of most transcripts coding for SCN-, ANO1- and TRPM-like channels are highest in head region of *Hydra*, including the hypostome and tentacles, as revealed by real-time PCR analysis. Mean+S.E.M., *n*=3–6. **g–h**, *In situ* hybridization with two probes specific for the transcripts *cluster2505* (coding for an ANO1-like channel) and *cluster30856* (SCN-like), confirms their localized expression in the base of tentacles. **i–m**, Immunohistochemical analysis using specific antibodies raised against synthetic peptides (confirmed the presence of ANO1-like (on i and l) and SCN-like (on j and m) channels in the neurons of the subpopulation N2 at the base of tentacles. No signal can be detected in the peduncle region of *Hydra* (on k). **n**, Taken together, the expression of gut dysmotility-associated ion channels identifies the nerve cell population N2 resident at the base of tentacles as putative pacemakers in *Hydra*. Scale bars: 100 µm (on g and h), 50 µm (on i, j, k) and 20 µm (on l and m). **Pharmacological interference experiments corroborate the essential role of Ano-, SCN- and TRPM-like channels and nicotinic acetylcholine receptors expressed in the neuronal population N2 in pacemaker activity in *Hydra*. o**, *Hydra* demonstrates a spontaneous rhythmic contractions followed by body extensions that occur on average every five minutes. **p**, The contraction pattern was video-recorded, transformed into a diagram of polyp shape, and used to assess the contraction frequency, defined as the number of full body contractions (red arrowheads) occurred within one hour, and the time intervals between consecutive contractions. **q**, Polyps were exposed to chemicals specifically modulating the activity of the ANO1-, SCN- and TRPM-like ion channels and nicotinic acetylcholine receptors (nAChR) expressed in the neuronal population N2. Experimental design: normal *Hydra* polyps were incubated in the inhibitors for 12 hours prior to a 1 hour recording of contractile behavior. Polyps incubated in 0.16% DMSO-supplemented (for Ani9) or pure (all other chemicals) *Hydra*-medium served as control. **r**, Contraction frequency is reduced in the presence of all chemicals targeting the channels expressed on the pacemaker population N2, but not affected in the presence of muscimol, which likely interferes with the population N7. **s**, Modulating the activity of the pacemaker-specific ANO1-, SCN- and TRPM-like channels and nAChRs in *Hydra* also disturbs the rhythmicity of spontaneous contractions, since the intervals between contractions become longer and less regular. Sampling size: *n*=10–49 animals (contraction frequency), *n*=25–283 intervals (interval length), * - p<0.05; ** - p<0.005; *** - p<0.0005; n.s. – p>0.05.

## ANO1-, SCN- and TRPM-like channels are essential for pacemaker activity in *Hydra*

We next tested the role of the neuronal subpopulation N2-specific ANO1-, SCN- and TRPM-like channels in controlling the pacemaker-driven rhythmic spontaneous contractions in *Hydra* (Fig. 2o–q). Exposing polyps to Ani9, a potent inhibitor of ANO1 channels^32^, resulted in both a two-fold reduction of the contraction frequency (Fig. 2q, r) and also in less regular contractions compared to controls (Fig. 2s). Similar results were obtained upon treatment of polyps with menthol, which activates TRPM8 channels in vertebrates^33^, and lidocaine, which interferes with SCN-like ion channels (Fig. 2q–s). The results show that modulating the activity of the neuron-specific ANO1-, SCN- and TRPM-like channels in *Hydra* greatly disturbs the rhythmicity of spontaneous contractions. Since neuronal subpopulation N2 is also characterized by the expression of putative nAChRs (Fig. 1g, Extended Data Fig. 3), we next tested the effects of tubocurarine (DTC), known as potent antagonist of nAChRs^34,35^, on the frequency and rhythmicity of the spontaneous contractions in *Hydra* (Fig. 2q). We found that the presence of DTC strongly reduced the frequency and affected the regularity of the spontaneous contractions (Fig. 2r, s). Other reflexes dependent on neural circuits such as the feeding reflex^36^ were not affected by most of the channel-specific inhibitors (Extended Data Fig. 12) indicating a specific role of the ion channels expressed in the N2 neurons in controlling rhythmic body contractions. Inhibiting GABAA receptors specifically expressed in neurons of subpopulation N7 and absent in the subpopulation N2 (Fig. 1g, Fig. 2q, Extended Data Fig. 3) had no effect onto the contraction pattern (Fig. 2r,s) but strongly influenced the feeding response (Extended Data Fig. 12). Together with the previous pharmacological findings^36^, our observations unequivocally identify the pacemaker population N2 as cholinergic, and the neuronal population N7 controlling the feeding response as predominantly GABAergic. Notably, all cells within neuronal population N2 homogeneously expressed high levels of one of the *innexin* genes – *cluster41630* (Extended Data Fig. 13). Innexins are the only components of gap junction complexes known in *Hydra*^37,38^. Homotypic gap junctions established between neurons of the N2 subpopulation therefore may electrically couple these neurons and allow generation of a neural network with pacemaker properties. In a similar way, a net of neurons present in the peduncle of *Hydra* is likely coupled by gap junctions^39^, and the epithelial cells of the bot, ectoderm and endoderm, are electrically coupled by gap junctions^40–42^. In sum, these data suggest that neurons of the N2 subpopulation express marker genes for gut dysmotility (Fig. 2a–n) and that inhibition of these ion channels disturbs spontaneous and regular body contractions (Fig. 2o–s). Therefore, neurons of the N2 subpopulation may act as pacemaker cells controlling the spontaneous contraction pattern. This is consistent with earlier electrophysiological recordings in *Hydra*^30^ and the observations that removal of the head region results in loss of spontaneous contractile activity^28^.

## Pacemaker neurons in *Hydra* are immunocompetent cells

We have shown previously that neurons in *Hydra* secrete antimicrobial peptides to shape the resident microbiome^43^. Neurons express a rich repertoire of peptides including the previously characterized antimicrobial peptide NDA1 and the dual-function neuropeptides RFamide III, Hym-370 and Hym-357 which previously were found to have strong activity against Gram-positive bacteria^43^ (Fig. 3a). In addition, neurons express homologs of Kazal2 and Arminin proteins that have been previously characterized^44–46^ as antimicrobial peptides in epithelial or gland cells in *Hydra* (Fig. 3a). Notably, the N2 pacemaker population also express multiple AMP molecules (Fig. 3a) indicating that these neurons, in addition to governing peristaltic motion, also are directly interacting with the resident microbiota. To determine if interaction with bacteria is a general characteristic of neurons in *Hydra*, we analyzed the seven neuronal subpopulations for expression of immune-related genes (Fig. 3b–d). Neurons express virtually all components of the Toll-like receptor (TLR/MyD88) pathway^47^ (Fig. 3b, Extended Data Fig. 14a) as well as many intracellular NACHT- and NB-ARC-domain containing NOD-like receptors^48^ (Fig. 3c, Extended Data Fig. 14b) and also C-type lectin (CTL) receptors (Fig. 3c, Extended Data Fig. 14c). Overall, these observations indicate that neurons in *Hydra* are immunocompetent cells, equipped with receptors, signal transducers and effector molecules to interact with bacteria.

**Fig. 3.**
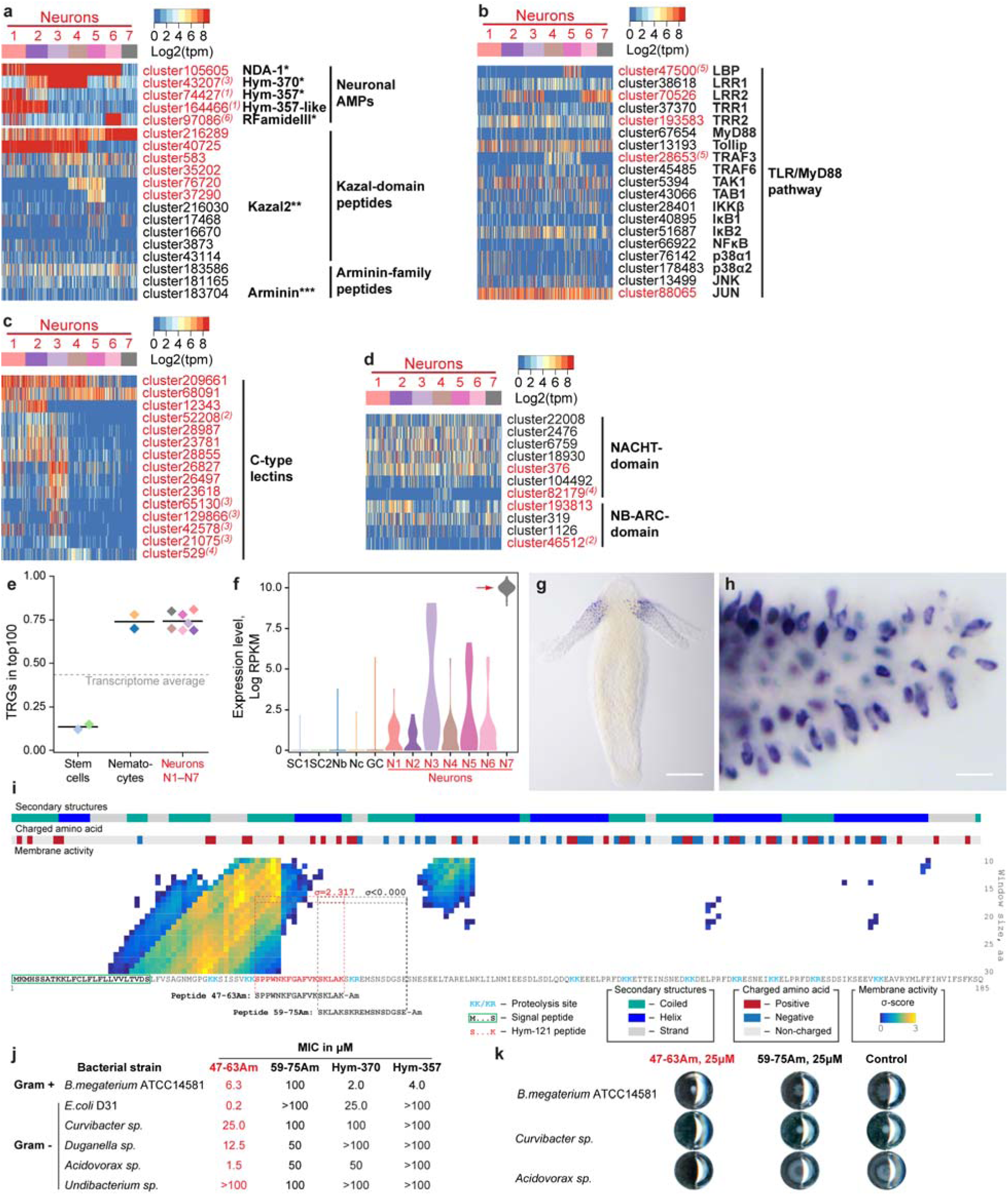
Neurons in *Hydra* are immunocompetent cells. **a**, Neurons express a rich set of peptides that have been previously characterized as antimicrobial peptides or their homologues. *- ref. ^43^; **- ref.^44^; ***- ref.^46^. **b**, Heatmap illustrates expression of transcripts coding for components of the Toll-like receptor (TLR)/MyD88-dependent immune pathway. Most components are present in the neurons, and five of them are significantly enriched in the neuronal population (red). Superscript numbers indicate the nerve cell cluster (N1-N7) where the transcripts are significantly (padj<0.05) enriched. **c**, Heatmap illustrates expression of some transcripts coding for NACHT- and NB-ARC-domain containing NOD-like receptors that have immune function. **d**, Multiple C-type lectin receptors that might recognize bacterial products are strongly expressed in the neurons. **e**, Over 70% of top100 transcripts specifically expressed in each of seven neuronal subpopulations (N1-N7) is represented by genes that have no homologues outside of Cnidaria, and thus are considered as TRGs. On the contrary, among the top 100 transcripts specifically enriched in the interstitial stem cells, only 15% are identified as TRGs. **f**, Transcripts coded by a TRG *cluster62692* are strongly upregulated in the neuronal subpopulation N7, weakly expressed in other neurons, and absent from non-neuronal cells of the interstitial lineage. **g–h**, *In situ* hybridization provides evidence that the TRG *cluster62692* is expressed exclusively in the sensory neurons of the tentacles. Scale bar: 100 µm (on g), 10 µm (on h). **i**, Moving window small-peptide scan prediction map for the peptide encoded by TRG *cluster62692* with residue charge and secondary structure annotations. The heat map reflects the peptide’s probability (σ-score) of being membrane active as predicted by the machine learning classifier^49^, and it is labeled at the first aa in the moving window frame. High σ-scores (yellow) suggest that *cluster62692* peptide is a potent antimicrobial peptide. An N-terminal signal peptide and multiple putative proteolysis sites, as well as a sequence identical to a previously described peptide Hym-121 (SPPWNKFGAFVKSKLAK-amide)^50^ are found within the *cluster62692* peptide, providing evidence that a preprohormone *cluster62692* is processed and gives rise to a secreted active peptide. The 17 aa long peptide corresponding to aa 47-63 (SPPWNKFGAFVKSKLAK=Hym-121) with high membrane activity score (σ=2.317) and control peptide aa 59-75 (SKLAKSKREMSNSDGSE) with no membrane activity score (σ=– 1.878) were synthesized, C-terminally amidated, and tested for antimicrobial activity in a Minimum Inhibitory Concentration (MIC) assay. **j**, The peptide 47-63Am is a potent antimicrobial peptide that shows selective growth inhibiting activity against Gram-positive and - negative bacteria. The control peptide 59-75Am demonstrates no antimicrobial activity. Consistently with the previous observations^43^, dual-function neuropeptides Hym-370 and Hym-357 show some antibacterial activity, yet weaker and more restricted than the peptide 47-63Am. **k**, Representative wells from plates of MIC assay. At concentration 25 µM, the peptide 47-63Am inhibits growth of *E. coli* and *Acidovorax sp*. and affects colony morphology of *B. megaterium*. The growth in the presence of control peptide (59-75Am, 25 µM) is not different from that in the pure medium (control).

In addition to a conserved toolbox of immune genes (Fig. 3b–d), *Hydra* neurons also employ some of their non-conserved, taxonomically-restricted genes (TRGs) to interact with bacteria. Non-conserved genes comprise over 70% of the cell-type specific genes in transcriptomes of the seven neuronal subpopulations (N1-N7) (Fig. 3e, Extended Data Fig. 15, Supplementary Data 2). The majority of the neuron-specific TRGs code for short peptides (<200 aa) with a N-terminal signal peptide sequence, but with no detectable structural domains (Supplementary Data 2). A machine learning-based approach^49^ identified that some of these novel genes code for putative peptides with high membrane destabilizing activity indicating strong antimicrobial activity (Extended Data Fig. 16, Supplementary Data 3). Surprisingly, the neuronal cluster N2 that contains the pacemakers was one of the populations most enriched in secreted peptides with putative antimicrobial activity (Extended Data Fig. 16). To provide direct evidence for functional relevance of TRGs in immune reactions in neurons, we characterized TRG *cluster62692* that is specifically expressed in neuronal subpopulation N7 in the tentacles (Fig. 3f–h) and encodes a 185 amino acid long peptide (Fig. 3i). The N-terminal signaling peptide is followed by a stretch of mostly positively-charged residues which is predicted to have a high membrane destabilizing activity (Fig. 3i). Screening a *Hydra* peptide database^50^ uncovered a 17 aa amidated peptide referred to as Hym-121 that is identical to aa 47-63 within the *cluster62692* polypeptide (Fig. 3i), thus providing an evidence for translation and proteolytic processing of the preprohormone encoded by TRG *cluster62692*. To test whether this peptide may serve as an antimicrobial peptide, we synthetized the amidated aa 47-63 peptide and subjected it to MIC assay against diverse bacteria. We found the peptide to inhibit growth of *Escherichia coli* at concentrations as low as 0.2 μM (Fig. 3j, k). The peptide also displayed a strong growth inhibiting activity (MIC 1.5 μM) against *Acidovorax* - a Gram-negative commensal microbe found on *Hydra*^51^. The growth of other two resident microbes, *Curvibacter* and *Duganella*, was impaired only at peptide concentrations 12.5-25 μM. Finally, the growth of *Undibacterium* was not inhibited by the 47-63 peptide even at 100 μM, indicating a high selectivity of the peptide’s antimicrobial activity (Fig. 3j). Surprisingly, while the peptide did not inhibit the growth of *Bacillus megaterium*, it drastically affected the colony morphology at concentrations as low as 6.3 μM (Fig. 3j, k). This suggests that the peptide modulates the motility, swarming behavior or spore formation of *B. megaterium*, implying thus a novel mechanism of AMP activity. Taken together, our data suggest that in addition to a conserved toolbox of immune genes (Fig. 3b–d), neurons also employ some of their TRGs to interact with bacteria supporting the view^52^ that the nervous system, with its rich repertoire of neuropeptides, is involved in controlling resident beneficial microbes.

## Murine intestinal pacemaker cells also are immunocompetent cells

To search for commonalities between *Hydra* and mice pacemaker neurons, we screened the transcriptome of murine intestinal pacemakers, the interstitial cells of Cajal (ICCs)^15^ for the presence of transcripts coding for immune receptors and pathways. Surprisingly, almost the entire signal-transducing cascade of the TLR/MyD88 pathway is present in the ICCs (Extended Data Fig. 17, Supplementary Data 4). ICCs also express antimicrobial peptides such as Defensin-8 (Supplementary Data 3) as well as components of the NLR- and CTL- pathways including NOD1, NOD2, and multiple CTL-domain receptors which are essential for detecting bacterial (Extended Data Fig. 18 and 19, Supplementary Data 4). This strongly suggests that ICCs in mice, similar to pacemaker neurons resident in subpopulation N2 in *Hydra*, are capable of interacting with bacteria and that, therefore, the molecular architecture of pacemaker cells is highly conserved in evolution (Fig. 4).

**Fig. 4.**
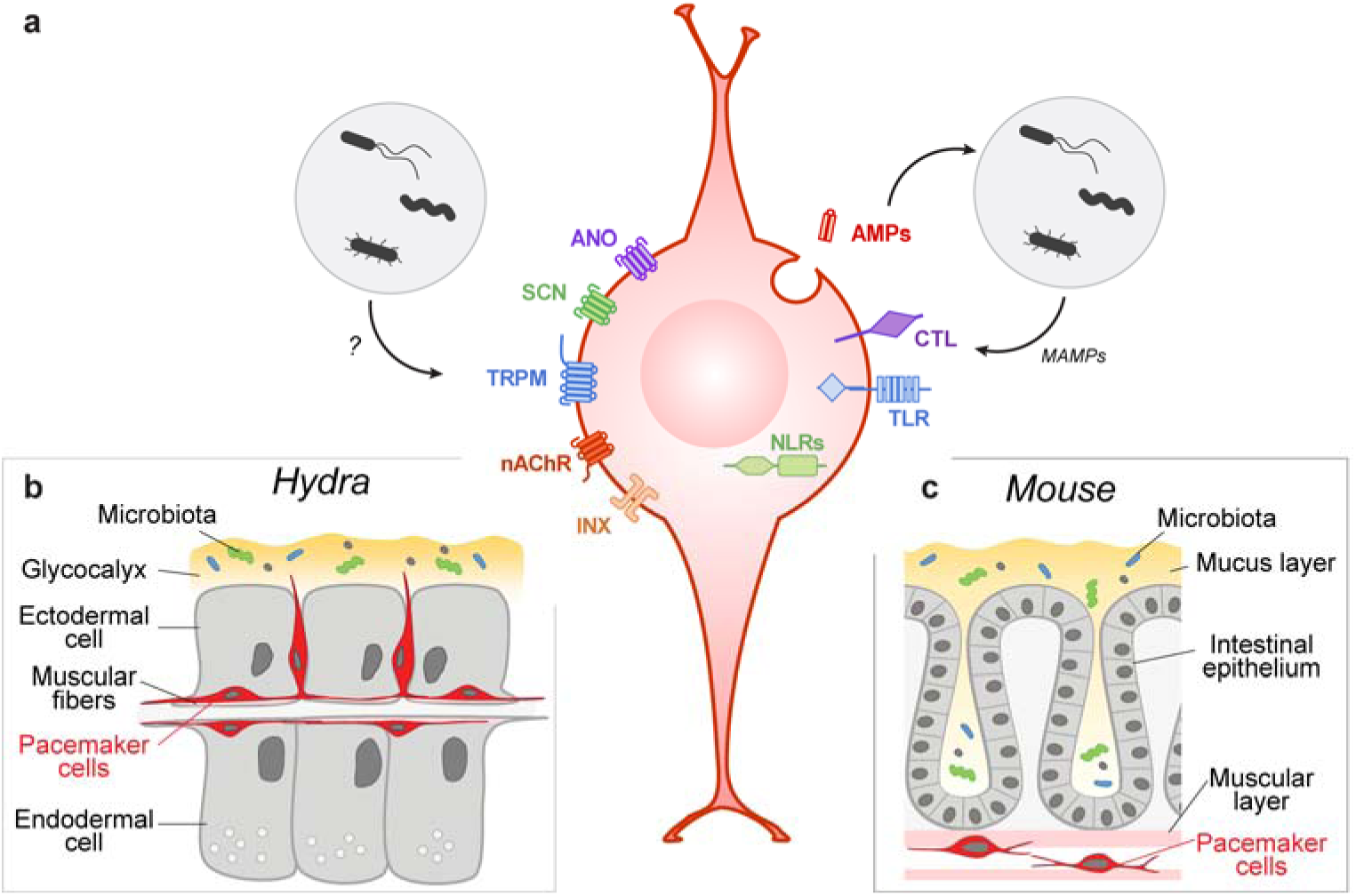
The molecular anatomy of the prototypical pacemaker cell. **a–c**, The gene expression program characteristic for pacemaker cells is highly conserved and is present in both, the neuronal population N2 that controls spontaneous contractions in *Hydra* (on b), and in the ICC driving the gut motility in mammals (on c). This evolutionary conserved signature of a pacemaker cell is made up of Ano-, SCN- and TRPM-like ion channels, nACh receptors and innexin gap junction. It also includes receptors such as Toll- and NOD-like receptors and C-type lectins, that are capable of recognizing bacterial products (MAMPs) and thus mediating the interaction between the host and its microbiota. The ion channels and neurotransmitter receptors might be also targets for bacteria-derived products.

## Conclusions

We have generated a comprehensive molecular profile of the neural subpopulations in *Hydra* and proposed a novel conceptual framework for phenotypic diversification of one of the simplest nervous systems in the animal kingdom. This framework builds upon seven spatially and functionally segregated neural subpopulations allowing for the emergence of diverse and complex behaviors within a morphologically simple nerve net structure.

Our data establish the neurons of the N2 population as a major contributor to controlling the rhythmicity of spontaneous body contractions in *Hydra*. These prototypical pacemaker cells reside in the head region and specifically express ANO1, SCN5 and TRPM-like ion channels that characterize human gut pacemakers. Consistently, the experimental inhibition of these channels greatly disturbs peristalsis in *Hydra*, which is consistent with our finding that these neurons are coupled by gap junctions into a network exhibiting pacemaker activity. The high degree of gene expression program conservation between the *Hydra* pacemaker subpopulation N2 and murine pacemaker cells supports that peristaltic motor activity of the gut is an evolutionary ancient archetypical property necessary to sustain life^53,54^, and that the cells with recurrent spontaneous electric activity have evidently emerged as early as the nervous system itself.

Finally, we discovered that *Hydra* pacemaker cells express a rich set of immune genes including antimicrobial peptides providing a mechanism for direct interference with resident microbes. The finding that many neuron-specific novel genes also encode antimicrobial peptides underlines the important function of neurons in interacting with microbes. Experimental interfering with microbiome in *Hydra*^26^ and disturbances of gut microbiota in humans^55–57^ similarly result in changes in pacemaker rhythmicity and abnormal peristalsis. Because the *Hydra* pacemaker neurons can directly mediate the interaction with microbiome, it might be plausible that human pacemakers in the gut similarly communicate with microbial communities. In fact, emerging data on mice provide first evidence for direct interactions between the gut microbiota, enteric neurons and the intestinal motility^58,59^. The evolutionary similarity or dissimilarity of the molecular toolkit used for such communication should become a subject of further investigations.

Altogether, our discoveries will improve the understanding of the archetypical properties of net nerve systems with pacemakers including human enteric nervous system, which is perturbed in human dysmotility-related conditions affecting a large portion of the general population worldwide. We therefore presume that the principles identified here are relevant far beyond *Hydra*.

## Methods

### Experimental design

Experiments were carried out using *Hydra vulgaris* strain AEP. Animals were maintained under constant conditions including the culture medium, food, and temperature (18°C) according to standard procedures^60^. Experimental animals were chosen randomly from clonally growing asexual *Hydra* cultures. The animals were typically fed three times a week, however they were not fed for 24 h prior to prior to pharmacological interference experiments, or for 48 h prior to RNA isolation, immunohistochemical staining and *in situ* hybridization.

### Generation of transgenic *Hydra* strains

To facilitate the FACS-mediated molecular profiling of single cells, we developed a transgenic *Hydra* line in which expressing two reporter proteins. The enhanced green fluorescent protein (eGFP) was cloned downstream from the previously reported *cnnos1* promoter^61^ and flanked by the actin terminator from the 3’-end. The second reporter, codon-optimized DsRED2 protein, was driven by the *actin* promoter sequence and flanked by the *actin* terminator region. This double cassette was cloned into the LigAF vector^62^. The transgenic construct was propagated in the *Escherichia coli* DH5-alpha strain and microinjected into fertilized embryos of *H. vulgaris* strain AEP as previously described^60,62^. Initial founder mosaic transgenic animals were clonally propagated, screened and enriched for transgenic cells until all interstitial stem cells were transgenic. The transgenic animals show no developmental abnormalities and are maintained in the lab for over 5 years.

### FACS isolation of cells

To isolate cells of the interstitial stem cell lineage from the transgenic *Hydra* by FACS (Extended Data Fig. 1a), the polyps were disintegrated into a cell suspension as previously described^61^. Briefly, 100 hydras were treated for 1.5□h in 1□ml dissociation medium (KCl 3.6 mM, CaCl2 6.0 mM, MgSO4 1.2 mM, Na citrate 6.0 mM, Na pyruvate 6.0 mM, TES 12.54 mM, glucose 6.0 mM, pH 6.9) supplemented with 50 U/mL Pronase E (Serva, Cat. No. 33635) on an orbital shaker at 200 rpm at 18°C. The resulting cell suspension was filtered through a 100 µm mesh to eliminate remaining tissue clumps and centrifuged for 5 min at 1,000 g at 4°C, the supernatant was removed, and the cells were resuspended in 1.0 mL cold dissociation medium. Single-cell suspensions were sorted according to FSC, SSC, eGFP and RFP fluorescence using FacsAria II cell sorting system (BD Biosciences, San Jose, CA, USA) utilizing the 100 µm nozzle and sheath pressure 20 PSI at 4°C. Gating was performed first using FSC and SSC to eliminate cell debris and doublets, and then the singlets were gated by the intensity of green and red fluorescent signals. First, a small sample was collected in the “bulk-sort” mode, checked microscopically using phase contrast and fluorescent microscopy with eGFP and RFP filter sets, and re-analyzed to verify purity of sorted fractions. In all cases, purity of the collected cell populations exceeded 95%. Further, single cells were sorted directly into 382-well plates containing 2.0 µL of lysis buffer supplemented with Triton X-100 and RNase inhibitor, 1.0 µL oligo-dT30VN primer and 1.0 µL dNTP mix per well. Immediately after FACS isolation, sorted cells were centrifuged at 700 g for 1□min, and snap frozen on dry ice. In total, 1152 individual cells were harvested: 384 GFP^−^/RFP^+^ neurons, 384 GFP^+^/RFP^−^ stem cells, and 384 GFP*^low^*/RFP*^low^* cells.

### Smart-seq2 library preparation and scRNA-sequencing

To generate cDNA libraries from the isolated cells, previously described Smart-seq2 protocol^63^ was implemented with minor modifications. After RNA denaturation at 72°C for 3 min and reverse transcription reaction, 22 cycles of pre-amplification were carried out using IS PCR primers. PCR products were purified on Ampure XP beads and the quality of cDNA libraries was assessed on an Agilent high-sensitivity DNA chip. A successful library had an average size of 1 kb. After tagmentation using Illumina Nextera XT DNA kit and amplification of adapter-ligated fragments using index N5xx and N7xx primers for 12 cycles, the PCR fragments were again purified, their concentration and size distribution were estimated using Qubit and Agilent chips, respectively. The libraries were further diluted to 2 nM, pooled, and paired-end sequenced on Illumina HiSeq2500 instrument. Raw sequences and quality scores for all clusters were extracted using CASAVA software.

### Reference transcriptome and gene annotation

A reference transcriptome of *H. vulgaris* strain AEP (accession number SRP133389) was assembled from 25 cDNA libraries generated from whole non-transgenic polyps at different conditions sequenced using Illumina HiSeq2500 v4 platform and annotated as previously described^64^. To generate the subsets of putative neurotransmitter receptors, ion channels and transcription factors, we used SMART, Pfam domain and PANTHER family predictions generated by InterProScan. To generate the proliferation signature dataset for cell type annotation, we identified in *Hydra* homologues of 25 highly conserved genes coding for proteins involved in cell division control and collectively known as “proliferation gene expression signature”^65^ using BLAST (Supplementary Data 5). To get a deeper insight into the top 300 genes differentially expressed in each of 12 clusters, we performed additional annotation steps. First, we performed BLAST analysis of the longest detected ORF predicted within each transcript using BLAST+ tool. Initially, BLAST was performed for all the differentially expressed transcripts from each cluster, using Uniprot database. A more precise BLAST search was then performed on the top 300 DE genes from each cluster, using NCBI nr Refseq database. Further, in order to detect TRGs among the DE genes, the BLAST search was performed first using the whole RefSeq database, and then with Cnidarian taxon (taxid 6073) excluded. A gene was considered as TRG if e-value from BLAST without cnidarian taxon is higher than 10e-10 and e-value from BLAST with cnidarian taxon is lower than 10e-10. Further, to predict the function of the putative gene products, we screened the predicted peptide sequences for the presence of signal peptides using SignalP^66^, the distribution of positively and negatively charged amino acids as well as putative strand, coiled or helical domains using GOR IV^67^ and inferred putative cellular localization using DeepLoc^68^. The sequences were also screened for the presence of putative structural domains using SMART^69^. Finally, to test, whether the peptides encoded within TRGs may possess properties of putative AMPs, the sequences were scanned for probability of having membrane disruption activity (σ-score) using a machine learning classifier^49^ with a moving window of 20 amino acids. The prediction scores are based on the support vector machine (SVM) distance to margin score, σ. Peptides with the membrane activity score σ>0.0 were considered as putative AMPs, and peptides with σ >1.0 – as high-confidence AMPs. To refine the prediction of an active peptide encoded within the TRG *cluster62692* precursor, we repeated the membrane activity scanning for this precursor using moving windows 10 to 30 aa. The results of prediction were visualized using heat maps. Supplementary Data 3 provides a list of all the TRGs screened using this machine learning tool with corresponding σ-score values.

### scRNA-seq data processing and quality control

Raw data from scRNA-seq were processed using a snakemake single cell RNA-seq pipeline in transcriptome mode. Briefly, data was mapped to the *H. vulgaris* transcriptome^64^ using STAR v2.5.2b^70^ with stringent gap penalties (*i.e*. outFilterMultimapNmax 100, alignIntronMax 1, alignIntronMin 2, scoreDelOpen −1000, scoreInsOpen −10000). Quality control was performed with the RSeQC package^71^ and gene expression was estimated with RSEM^72^ with groups of contigs (called clusters) defined as genes as previously described^64^. Low quality cells were filtered according to the following criteria: < 20% uniquely mapping reads, > 26.2% ERCC spike-in mapping, >68.7% rRNA mapping, >13.9% of the reads at the 10% most 3’ end of transcripts (using only transcripts with complete ORFs) and < 3,000 genes detected, < 798,442 counts mapped. Any cell that failed two of these criteria was removed from the analysis.

### scRNA-seq analysis

Cells passing quality control (*i.e*. 1,016) were analyzed using the R package Seurat_2.3.4. We used specific parameters for the different input gene sets. Input genes for each gene set are in Supplementary Data 1, 7 and 8.

### Whole transcriptome

We included genes detected in at least 3 cells and cells that contained at least 5,000 transcripts. This resulted in a total of 928 cells and 166,186 transcripts. Subsequently, we performed normalization using the option “LogNormalize” with scale.factor the mean number of genes per cell. We selected a total of 6,549 variable genes (x.low.cutoff=0.5, x.high.cutoff=50, y.cutoff=0.5) and identified cell clusters using 20 PCA dimensions and a resolution parameter equal to 0.6.

### Receptors

We included genes detected in at least 1 cells and cells that contained at least 1 transcript in order to include all receptor genes. This resulted in a total of 926 cells and 112 transcripts (Supplementary Data 7). Subsequently, we performed normalization using the option “LogNormalize” with scale.factor the mean number of genes per cell. We used the 112 receptor genes as variable genes (x.low.cutoff=0, x.high.cutoff=7, y.cutoff=−2) and identified cell clusters using 6 PCA dimensions and a resolution parameter equal to 0.6.

### Ion channels

We included genes detected in at least 2 cells and cells that contained at least 2 transcripts in order to include all ion channel genes. This resulted in a total of 928 cells and 431 transcripts (Supplementary Data 8). Subsequently, we performed normalization using the option “LogNormalize” with scale.factor the mean number of genes per cell. We used the 431 ion channel genes as variable genes (x.low.cutoff=0, x.high.cutoff=50, y.cutoff=−3) and identified cell clusters using 15 PCA dimensions and a resolution parameter equal to 0.6.

### Transcription factors

We included genes detected in at least 1 cells and cells that contained at least 1 transcript in order to include all transcription factor genes. This resulted in a total of 928 cells and 364 transcripts (Supplementary Data 1). Subsequently, we performed normalization using the option “LogNormalize” with scale.factor the mean number of genes per cell. We used the 364 ion channel genes as variable genes (x.low.cutoff=0, x.high.cutoff=7, y.cutoff=−3) and identified cell clusters using 5 PCA dimensions and a resolution parameter equal to 0.6.

### Hierarchical clustering

The input data was TPM normalized in RSEM and subset for the corresponding input cells and genes that underwent dimensionality reduction. A +1 pseudocount was added prior to log2 transformation. Heatmaps were generated in R using the heatmap.2 function based on Euclidean distances and the ward.D2 method.

### *In situ* hybridization

To map the seven neuronal clusters populations in the *Hydra* body, we performed *in situ* hybridization with a set of genes strongly enriched in either of the seven neuronal subpopulations. Expression patterns were detected in the whole mount *Hydra* preparations by *in situ* hybridization with an anti-sense digoxigenin-labeled RNA probes as previously described^73^. DIG-labeled sense-probe was used as a control. Signal was developed using anti-DIG antibodies conjugated to alkaline phosphatase (1:2000, Roche) and NBT/BCIP staining solution (Roche). Images of the *in situ* preparations were captured on a Zeiss Axioscope with Axiocam camera.

### Pharmacological interference assays

To investigate the role of specific ANO1-, SCN- and TRPM-like channels in the pacemaker activity in *Hydra*, we exposed normal *H. vulgaris* AEP polyps to different pharmacological agents, recorded and quantified their behavior. Polyps were treated with 25 µM Ani9 (Sigma, Cat. No. SML1813), 200 µM menthol (Sigma, Cat. No. 15785), 100 µM lidocaine (Sigma, Cat. No. L5647), 1 mM tubocurarine (DTC, Sigma, Cat. No. 93750) or 100 µM muscimol (Sigma, Cat. No.M1523) for 1 or 12 hours at 18°C. Control polyps were incubated either in Hydra-medium or in the medium supplemented with 0.16% DMSO (for Ani9, which has been dissolved in 100% DMSO to stock concentration 15 mM). The spontaneous contractions were video-recorded and quantified as previously described^26^. We recorded the behavior for 90 min with a frequency 20 frames per minute. For further analysis, we excluded first 30 min of the recorded sequence, and quantified number of full body contractions and their periodicity using a custom ImageJ plugin^26^. The contraction frequencies were normalized to the average frequency of contractions in corresponding control polyps. To examine the effects of the modulators on the feeding reflex, *Hydra* polyps were pretreated with the pharmacological agents, their feeding reflex was elicited by 10 µM reduced glutathione (GSH, Sigma, Cat. No. G4251), and the duration of feeding response was recorded as described by Lenhoff^74^.

### Statistical analysis

The sample size (*n*) reported in the figure legends is the total amount of animals used in each treatment. Each animal employed was assigned to only one treatment and was recorded only once. Treatment of the polyps with pharmacological substances and evaluation of the behavioral parameters (contraction frequency, intervals between contractions, and duration of feeding response) was blinded. Differences in contraction frequency, interval between contractions, and feeding response duration between the treatments (*i.e*. Ani9, lidocaine, menthol, DTC, and muscimol) and corresponsing control (*i.e.* Hydra-medium or DMSO) were analyzed using unpaired *t*-test.

### MIC determination of antimicrobial activity

To test whether peptides encoded in TRGs may have an antimicrobial function, two 17 aa long peptides corresponding to aa 47-63 (SPPWNKFGAFVKSKLAK) and aa 59-75 (SKLAKSKREMSNSDGSE) of the prepropeptides encoded by the *cluster62692* were synthesized (up to 5 mg), C-terminally amidated and purified to a purity of >□95% (GenScript, USA), and their antimicrobial activity was estimated in a Minimum Inhibitory Concentration (MIC) assay as previously described^43^. The following bacterial strains were used in MIC assays: *Bacillus megaterium* ATCC14581, *Escherichia coli* D31, and four isolates from the natural *H. vulgaris* strain AEP microbiota: *Curvibacter sp*., *Duganella sp.*, *Acidovorax sp*., and *Undibacterium sp.*^46,75^. Microdilution susceptibility assays were carried out in 96-well microtiter plates that were pre-coated with sterile 0.1% bovine serum albumin (BSA). After removal of BSA the wells were filled with a twofold dilution series of either aa 47-63 or aa 59-75 peptide. We also tested the neuropeptides Hym-370 (KPNAYKGKLPIGLW-amide) and Hym-357 (KPAFLFKGYKP-amide) that has been previously identified as putative antimicrobial substances^43^. Lyophilized peptides were dissolved in ultrapure water to stock concentration of 10 mg/mL. Incubation with an inoculum of approximately 100 CFU per well was performed in PBS buffer (pH 6.2) overnight at 37□°C for *B. megaterium* and *E. coli*, or in R2A media for 3-4 days at 18°C for four isolates of *Hydra* bacteria. The MIC was determined as the lowest serial dilution showing absence of a bacterial cell pellet. Experiments were carried out in triplicates.

### Phylogenetic analysis of ion channel genes

To uncover, whether *Hydra* has homologues of the ion channel genes whose expression is known to be either restricted to mammalian ICCs or essential for gut motility, we performed a BLAST search (tblastn) using full-length amino acid sequences of ANO1 (UniProt accession number Q5XXA6), SCN5A (accession number Q14524) and TRPM8 (accession number Q7Z2W7) proteins from *Homo sapiens* against the genome of *H. vulgaris*^38^ (available at https://research.nhgri.nih.gov/hydra/) and the reference transcriptome of *H. vulgaris* strain AEP (SRA accession SRP133389). Matches with expectation e-value <10e-10 were considered as signs of homolog presence and were verified by manual domain composition analysis using SMART^69^, transmembrane domain prediction with TMHMM^76^ and reciprocal BLAST against the UniProt database. Maximum-likelihood phylogenetic trees of ANO1, SCN5A and TRPM8 homologs from *Hydra*, human, African clawed frog *Xenopus laevis*, and zebrafish *Danio rerio* were built using full-length amino acid sequences aligned using MUSCLE^77^ with 1000 bootstrap iterations.

### Quantitative real-time PCR gene expression analysis

To test, whether the genes coding for ANO1, SCN5A and TRPM8 homologues in *Hydra* are differentially expressed in the polyp along the oral-aboral axis, we performed quantitative real-time PCR. We dissected polyps into three body sections: head (hypostome area with tentacles), body column, and foot (peduncle). Each total RNA was extracted from body fragments obtained from 50 polyps, and converted into the cDNA as previously described^78^. Real-time PCR was performed using GoTaq qPCR Master Mix (Promega, USA) and oligonucleotide primers specifically designed to amplify the homologues of ANO1, SCN5A and TRPM8 ion channel genes, as well as the *ef1a* (translation elongation factor 1 alpha) and *actin* genes as equilibration references (Supplementary Table 1). For each body part, we made 3-5 biological replicates. The data were collected by ABI 7300 Real-Time PCR System (Applied Biosystems, USA) and analyzed by the conventional ΔΔCt method.

### Generation of antibodies and immunohistochemistry

To localize the expression of ANO1-like and SCN5A-like ion channels in *Hydra* using immunocytochemistry, polyclonal antibodies were raised against synthetic peptides in rabbits. The peptides that correspond to intracellular loops located between transmembrane domains of the ion channels (hySCN5-like: SRSKPKMFKDYKPE; hyANO-like: ETRRPIADRAQD) were synthetized, purified and N-terminally conjugated with KLH prior to injection (GenScript, USA). Polyclonal antibodies were affinity purified on the antigen and concentrated to 1.5 mg/mL. Serum harvested from the rabbits prior to their immunization was used as control.

Immunohistochemical detection in whole mount *Hydra* preparations was carried oud as described previously^73^. Briefly, polyps were relaxed in 2% urethane in Hydra-medium, fixed in 4% (v/v) paraformaldehyde, washed with 0.1% Tween in PBS, permeabilized with 0.5% Triton X-100 in PBS, incubated in blocking solution (1% BSA, 0.1% Tween in PBS) for 1 h and incubated further with primary antibodies diluted to 1.0 μg/mL in blocking solution at 4°C. AlexaFluor488-conjugated goat-anti-rabbit antibodies (Invitrogen, USA) were diluted to 4 μg/mL in blocking buffer and incubations were done for 1 h at room temperature. Rhodamin-phalloidin (Sigma) and TO-PRO3-iodide-AlexFluor633 (Invitrogen, USA) counterstaining was conducted as described previously^73^. Confocal laser scanning microscopy was done using a TCS SP1 laser scanning confocal microscope (Leica, Germany).

## Supplementary information is available for this paper

Extended Data Figures 1 to 19, Extended Data 1 to 8, Supplementary Table 1,

## Acknowledgements

We thank Vassilis Pachnis for critically reading the manuscript and for sharing unpublished observations in mouse enteric neurons. The authors are thankful to Eva Herbst (Kiel University), Hans-Heinrich Oberg (UKSH, Kiel), Kiran Sedimbi (SciLifeLab, Stockholm), Simone Picelli (IOB, Basel) and The Eukaryotic Single Cell Genomics (ESCG) facility at SciLifeLab for technical support, and to the Uppsala Multidisciplinary Center for Advanced Computational Science (UPPMAX) for providing computational infrastructure. The authors also appreciate the help of Jan Taubenheim (Heinrich Heine University, Düsseldorf) and Tatiana Chontorotzea (Medical University of Vienna) for help in analyzing the data. The authors are thankful to Toshitaka Fujisawa (COS, Heidelberg) for providing access to Hydra Peptide database.

## Author contributions

A.K., M.K., C.G., A.M., A.-S. M., G.C. acquired all biological data and performed the relevant analysis. S.G. analyzed single-cell data. Å.B. performed read alignment and quality filters. S.G., L.F. annotated differentially expressed genes. J.d.A. and G.C.L.W. developed and implemented computational methods. A.K., M.D’A., I.A., T.C.G.B. conceptualized and designed the study and wrote the manuscript. A.K., I.A. and T.C.G.B. supervised the study and acquired the financial support for the study.

## Data and materials availability

Raw sequence reads from single-cell RNA-seq datasets have been deposited in the GEO under accession code XXXXXX. The pipeline used for read mapping and quantification of scRNA-seq data is available at: https://bitbucket.org/scilifelab-lts/lts-workflows-sm-scrnaseq. R code written to analyze the scRNA-seq data is available upon request. The transgenic *Hydra* line is available from the laboratory of Thomas C.G. Bosch. All data needed to evaluate the conclusions in the paper are present in the paper or the supplementary materials.

