## Supplementary Information for "Prototypical pacemaker neurons are immunocompetent cells"

**This PDF file includes:**

Extended Data Fig. 1 to 19
Supplementary Table 1
Captions for Extended Data 1 to 8

**Other Supplementary Materials for this manuscript include the following:**

Extended Data 1 to 8

**Extended Data Fig. 1**

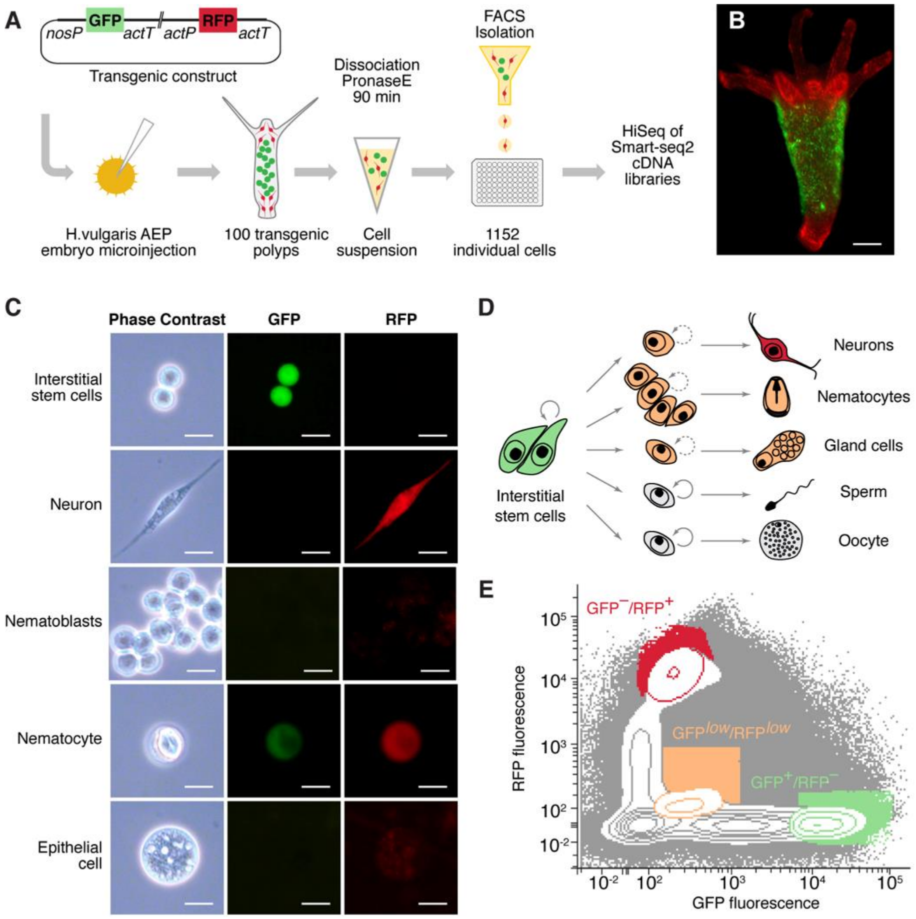


**Differential fluorescent labeling and FACS-mediated isolation of interstitial cell lineage.** (**A**) The workflow of the single-cell molecular profiling of *Hydra* interstitial cell lineage. (**B**) A transgenic polyp expressing GFP in the interstitial stem cells in the middle body compartment and RFP in the neurons concentrated in the tentacles, hypostome, and the foot region. (**C**) Phase contrast and fluorescent microscopy of disintegrated cells from transgenic *Hydra*. Interstitial stem cells are strongly GFP-positive but show now RFP fluorescence (GFP$^{+/}$RFP$^{-}$). Mature neurons are strongly RFP-positive, while demonstrating no GFP fluorescence (GFP$^{-}$/RFP$^{+}$). Nematoblasts and nematocytes showed weak GFP and weak RFP fluorescence (GFP$^{low}$/RFP$^{low}$). Epithelial cells demonstrate only weak unspecific red fluorescence that is microscopically visible but is not detected by FACS. Scale bar: 10 µm. (**D**) Combination of two fluorescent proteins allowed differential labeling of all somatic cell types within the interstitial cell lineage. Interstitial stem cells continuously proliferate and differentiate into neurons, gland cells, nematocytes, and gametes. (**E**) Three cell populations were isolated from dissociated transgenic *Hydra* polyps using FACS based on their relative GFP and RFP fluorescence: $GFP^{+}/RFP^{–}$, $GFP^{–}/RFP^{+}$, and $GFP^{low}/RFP^{low}$.

**Extended Data Fig. 2**

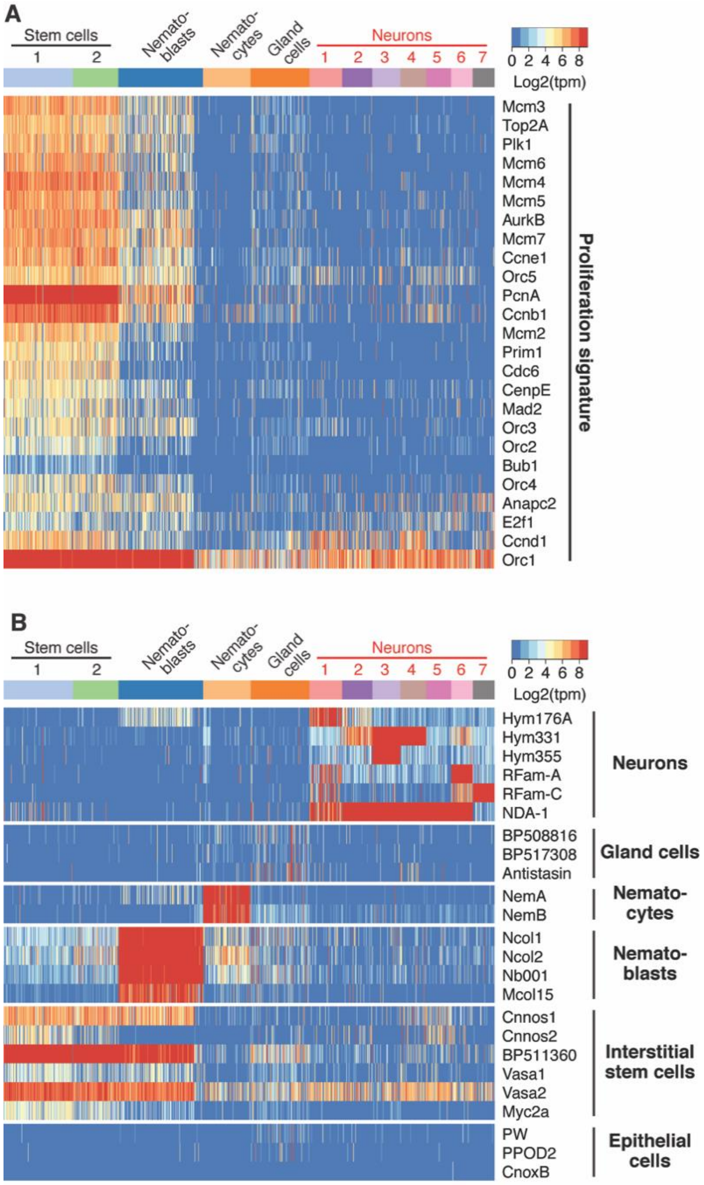


**Annotation of cell populations using the expression profiles of the proliferation signature and cell-type specific genes. (A)** Heatmap illustrating expression of genes comprising the proliferation signature (see Exteded Data 5). These genes are strongly expressed in proliferating stem cells and nematoblasts and are not expressed in the differentiated non-dividing neurons and nematocytes. (**B)** Heatmap illustrates expression of specific marker genes used to annotate the 12 clusters (see Extended Data 6). Markers specific for the interstitial stem cells and their progeny (neurons, nematoblasts, nematocytes, gland cells) demonstrate strong and distinct expression pattern. Marker genes characteristic for the epithelial cells are not expressed in the dataset, providing evidence for specific profiling of the interstitial lineage without contamination from the ectodermal and endodermal cells.

**Extended Data Fig. 3**

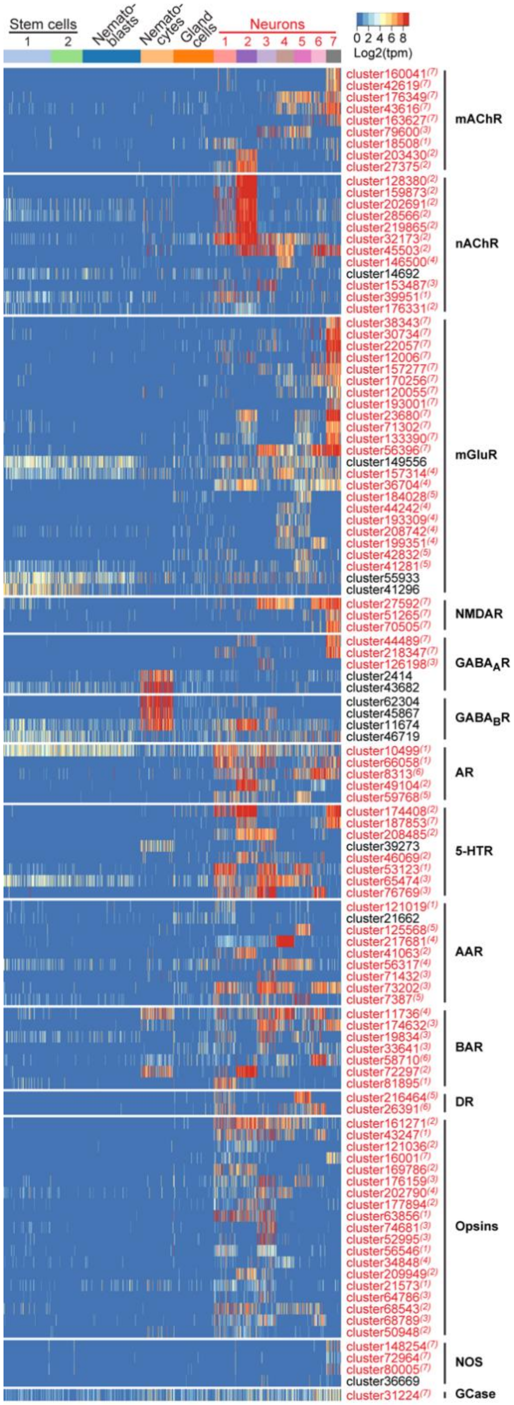

**Each neuronal population is characterized by a specific set of transcripts coding for receptors for neuromediators.** Heatmap illustrating expression levels of genes coding for putative neurotransmitter receptors (see Extended Data 7) in the interstitial stem cell lineage: muscarinic (mAChR) and nicotinic (nAChR) acetylcholine receptors, metabotropic (mGluR) and N-methyl-D-aspartate (NMDAR) glutamate receptors, A-type ($GABA_AR$) and B-type ($GABA_BR$) γ-aminobutyric acid, adenosine (AR) and serotonin (5-HTR) receptors, alpha- (AAR) and beta- (BAR) adrenergic receptors, dopamine receptor (DR). Multiple transcripts coding for putative light-sensitive receptors of opsin family are present in the neurons of *Hydra*. Genes coding for nitric oxide synthases (NOS) are expressed exclusively in the neuronal population N7. Transcripts coding for the guanylate cyclase enzymes (GCase) that may serve as a receptor for NO secondary messenger are also enriched in the subpopulation N7. Transcripts specifically upregulated in the neurons are labelled red, superscript numbers indicate the nerve cell cluster (N1-N7) where the transcripts are significantly ($padj<0.05$) enriched.

**Extended Data Fig. 4**

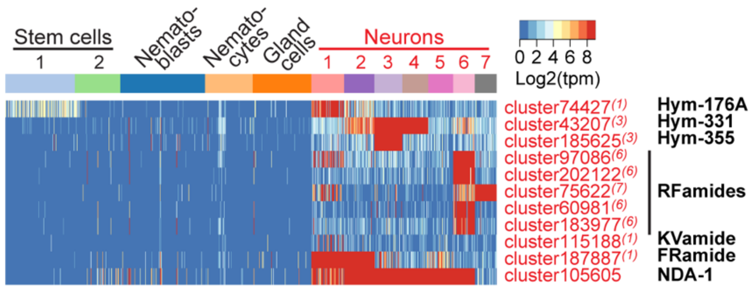


**Each neuronal population is characterized by a specific set of transcripts coding for neuropeptides.** Heatmap illustrating expression of genes coding for characterized[50] peptides Hym-176, Hym-331, and Hym-355, five paralogues of RFamide family, a KVamide, FRamide, and recently characterized neuron-derived antimicrobial peptide NDA-1[43]. Transcripts specifically upregulated in the neurons are labelled red, superscript numbers indicate the nerve cell population (N1-N7) where the transcripts are significantly (padj<0.05) enriched.

**Extended Data Fig. 5**

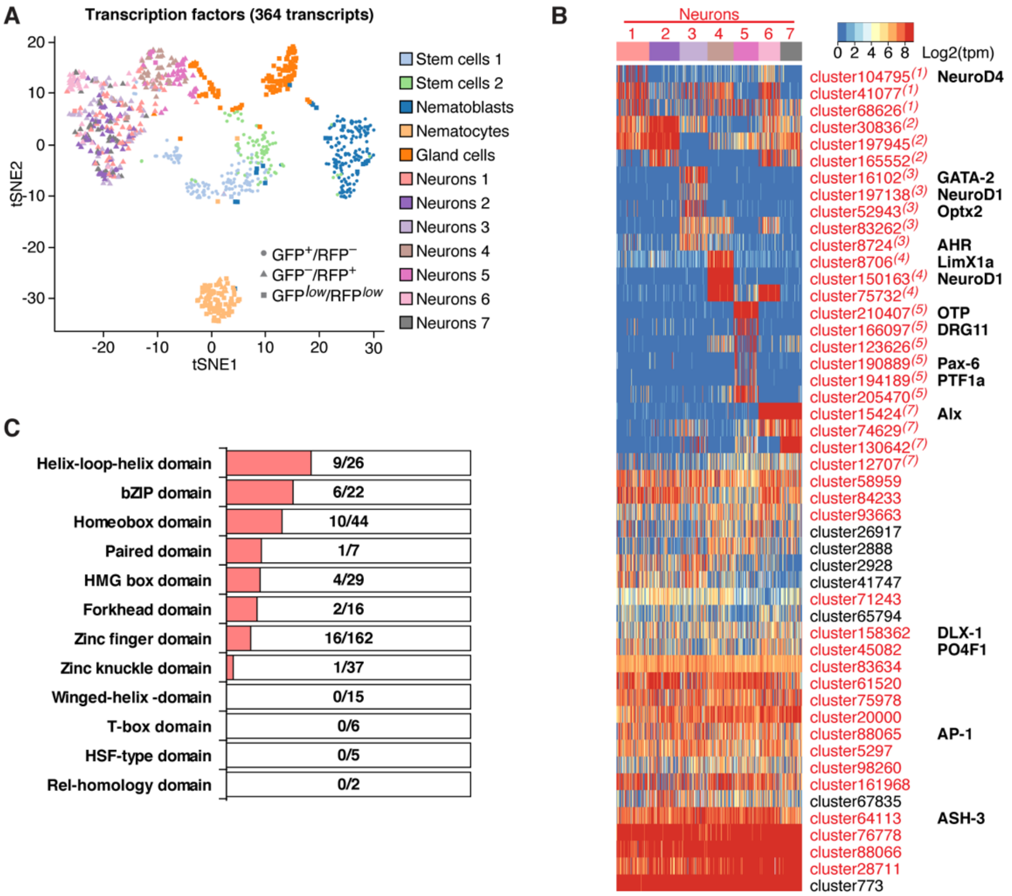


**Seven neuronal subpopulations express a common set of transcription factors**. (**A**) t-SNE map based on expression analysis of 364 transcripts coding for transcriptional factors (TFs, see Extended Data 1). All seven populations of neurons express common set of TFs and are not segregated on the t-SNE plot, in contrast to the stem cells, nematoblasts, nematocytes and the gland cells. (**B**) Heatmap illustrating expression levels of transcripts coding for TFs and highly abundant in the neurons. Transcripts specifically upregulated in the neurons are labelled red, superscript numbers indicate the nerve cell cluster (N1-N7) where the transcripts are significantly (padj<0.05) enriched. In addition to common neuron-specific TFs, each neuronal population is characterized by a combinatorial expression of few genes encoding other TF, likely acting as selector genes.

Notably, multiple homologues of neurogenic TFs found in Bilateria (right column) are present in distinct neuronal populations, suggesting their conserved role in neurogenesis in the animal kingdom. (**C**) The neuron-specific TF signature is composed of mainly of Zn-finger, homeodomain and helix-loop-helix DNA-binding proteins. On the contrary, members of other families, such as Winged-helix, T-box, and Rel-homology domain TFs, are absent from the neurons.

**Extended Data Fig. 6**

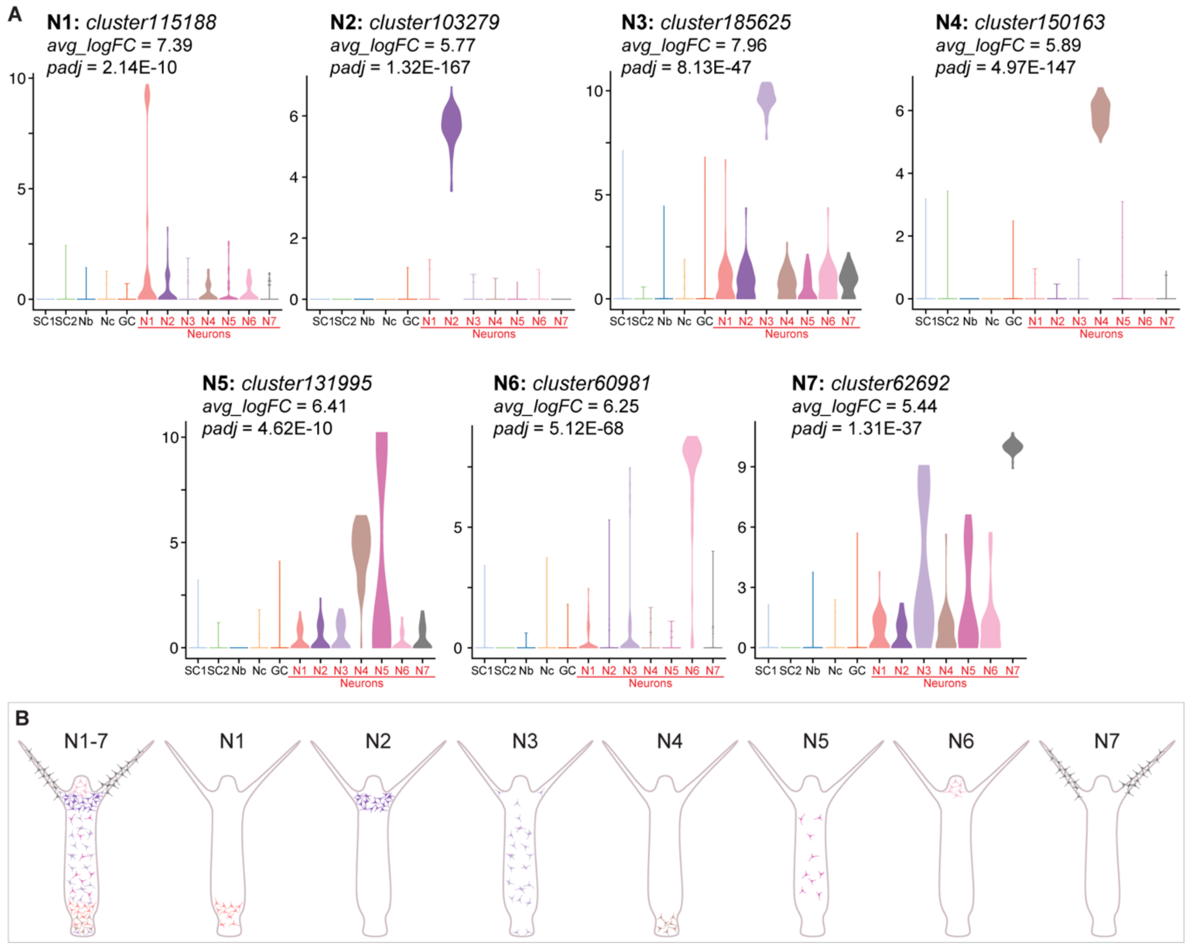


**Expression of marker genes strongly enriched specific nerve cells subpopulations (N1-N7) used to map the neuronal populations in *Hydra*.** (**A**) Violin plots demonstrating expression levels of seven marker genes used to map distinct neuronal populations by *in situ* hybridization (Fig. 1j) in all cells within the interstitial cell lineage. Y-axis represents expression level as LogRPKM. Average upregulation fold change (avg_logFC) and adjusted p-value (padj) are indicated for each gene. (**B**) The nerve net of *Hydra* is organized in seven spatially restricted cell populations. Localization of the populations is inferred from the *in situ* hybridization results (Fig. 1j).

**Extended Data Fig. 7**

a

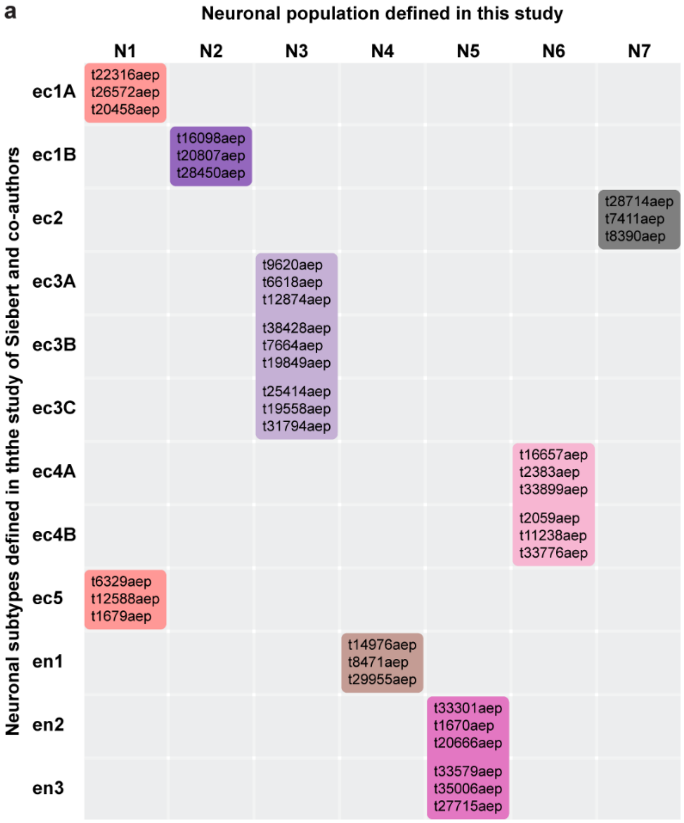


b

| Transcript ID, Siebert et al. | Transcript ID, our study | Cluster, our study | logFC | Padj |
|---|---|---|---|---|
| t22316aep | cluster60609 | N1 | 5.13 | 3.33E-49 |
| t26572aep | cluster33557 | N1 | 3.16 | 1.30E-32 |
| t20458aep | cluster200791 | N1 | 3.08 | 1.05E-10 |
| t16098aep | cluster103279 | N2 | 5.77 | 1.32E-167 |
| t20807aep | cluster66153 | N2 | 4.43 | 1.40E-84 |
| t28450aep | cluster119726 | N2 | 5.53 | 2.52E-88 |
| t28714aep | cluster56664 | N7 | 2.72 | 2.53E-86 |
| t7411aep | cluster94531 | N7 | 8.77 | 5.72E-38 |
| t8390aep | cluster62692 | N7 | 5.44 | 1.31E-37 |
| t9620aep | cluster199921 | N3 | 3.10 | 1.48E-42 |
| t6618aep | cluster124097 | N3 | 1.05 | 6.07E-38 |
| t12874aep | cluster185625 | N3 | 7.96 | 8.13E-47 |
| t38428aep | cluster52943 | N3 | 5.33 | 9.33E-58 |
| t7664aep | cluster160659 | N3 | 7.20 | 1.79E-46 |
| t19849aep | cluster155448 | N3 | 4.31 | 2.19E-26 |
| t25414aep | cluster65043 | N3 | 3.54 | 1.85E-34 |
| t19558aep | cluster137620 | N3 | 4.77 | 8.58E-87 |
| t31794aep | cluster106527 | N3 | 5.75 | 1.42E-56 |
| t16657aep | cluster183977 | N6 | 8.18 | 1.68E-39 |
| t2383aep | cluster58169 | N6 | 5.72 | 2.69E-109 |
| t33899aep | cluster60981 | N6 | 6.25 | 5.12E-68 |
| t2059aep | cluster202122 | N6 | 8.34 | 6.69E-50 |
| t11238aep | cluster127881 | N6 | 1.54 | 5.79E-03 |
| t33776aep | cluster61788 | N6 | 2.79 | 4.78E-22 |
| t6329aep | cluster115188 | N1 | 7.39 | 2.14E-10 |
| t12588aep | cluster193988 | N1 | 6.77 | 4.91E-71 |
| t1679aep | cluster74427 | N1 | 7.60 | 1.12E-35 |
| t14976aep | cluster150163 | N4 | 5.89 | 4.97E-147 |
| t8471aep | cluster133757 | N4 | 8.11 | 1.00E-56 |
| t29955aep | cluster160146 | N4 | 5.94 | 2.31E-80 |
| t33301aep | cluster206126 | N5 | 7.03 | 3.66E-33 |
| t1670aep | cluster138649 | N5 | 7.43 | 1.42E-39 |
| t20666aep | cluster153425 | N5 | 7.09 | 2.85E-47 |
| t33579aep | cluster53954 | N5 | 7.27 | 6.79E-24 |
| t35006aep | cluster44629 | N5 | 6.35 | 8.80E-102 |
| t27715aep | cluster108398 | N5 | 5.02 | 1.49E-16 |

**Correspondence between 7 neuronal clusters identified in our study (N1-7) and 12 neuronal subtypes reported by Siebert and co-authors[27].** Expression analysis of marker genes specifically expressed in each of 12 neuronal subtypes reported by Siebert and co-authors (see Fig. S41 in ref. 27) uncovers a clear correspondence between the clusters. Marker genes characteristic for 12 neuronal subtypes from study of Siebert and co-authors have corresponding transcripts in our dataset that are significantly enriched in specific neuronal subpopulations (on B).

**Extended Data Fig. 8**

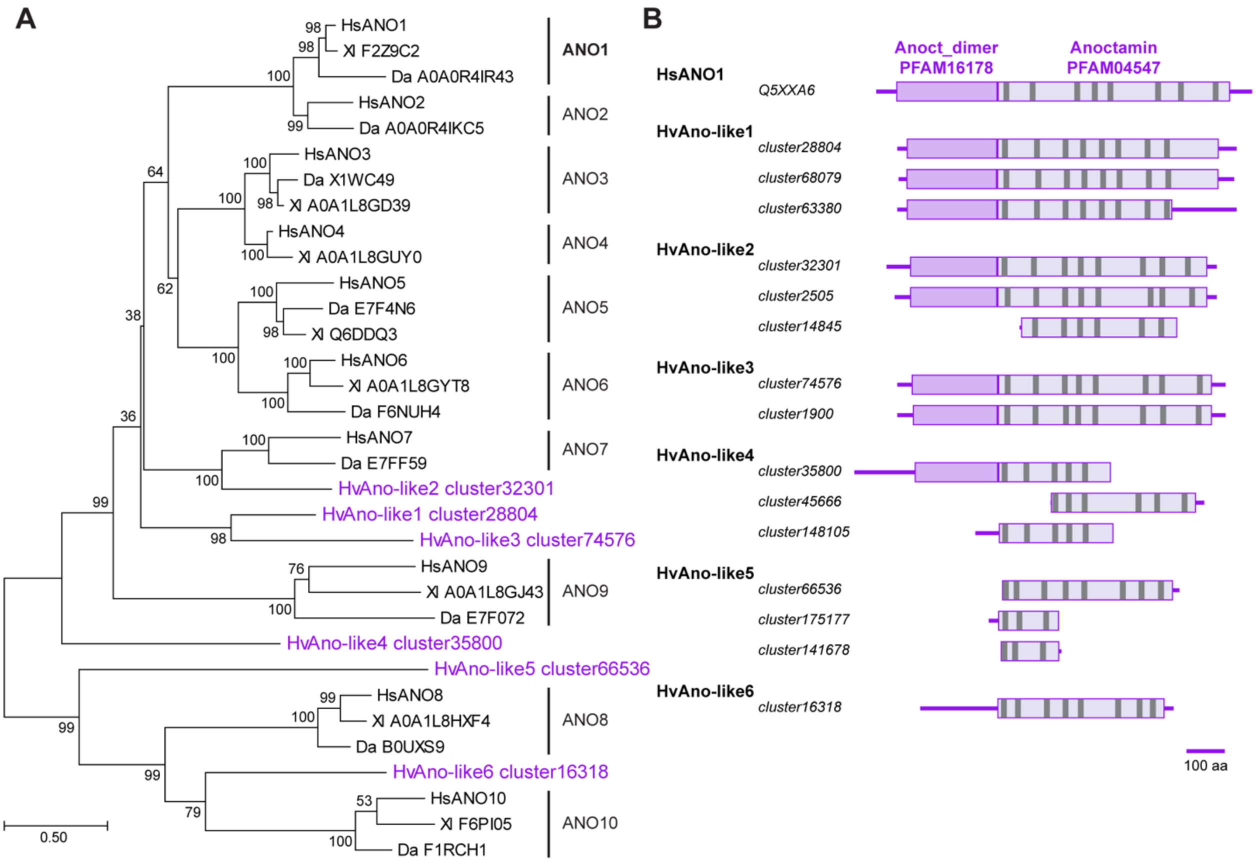


**Pacemaker-specific Ca-activated chlorine channels of Anoctamin family are highly conserved in *Hydra*.** (**A**) Detailed phylogenetic analysis of Anoctamin ion channels from human (*Homo sapiens*, Hs), frog (*Xenopus laevis*, Xl), zebrafish (*Danio rerio*, Da), and hydra (*Hydra vulgaris*, Hv). *Hydra* possesses six genes coding for anoctamin-like ion channels – HvAno-like1–6. The evolutionary history was inferred by using the Maximum Likelihood method based on the Le and Gascuel model[79]. The tree with the highest log likelihood is shown. The percentage of trees in which the associated taxa clustered together is shown next to the branches. The tree is drawn to scale, with branch lengths measured in the number of substitutions per site. Only fewer than 10% alignment gaps, missing data, and ambiguous bases were allowed at any position. There were a total of 549 positions in the final dataset. Evolutionary analyses were conducted in MEGA7[80]. (**B**)

Domain structure analysis using SMART and TMHMM algorithms uncovers remarkably high conservation in structure between the human ANO1 protein and six anoctamin-like channels from *Hydra*. Multiple transcripts corresponding to the same gene were frequently identified in the reference transcriptome of *Hydra*, suggesting either existence of alternative splice-variants or poor assembly of the full-length transcripts, most likely due to very low expression level of the genes (see Extended Data Fig. 11). Localization of specific Anoctamin dimer (PFAM16178) and Anoctamin (PFAM04547) domains and transmembrane domains is illustrated.

**Extended Data Fig. 9**

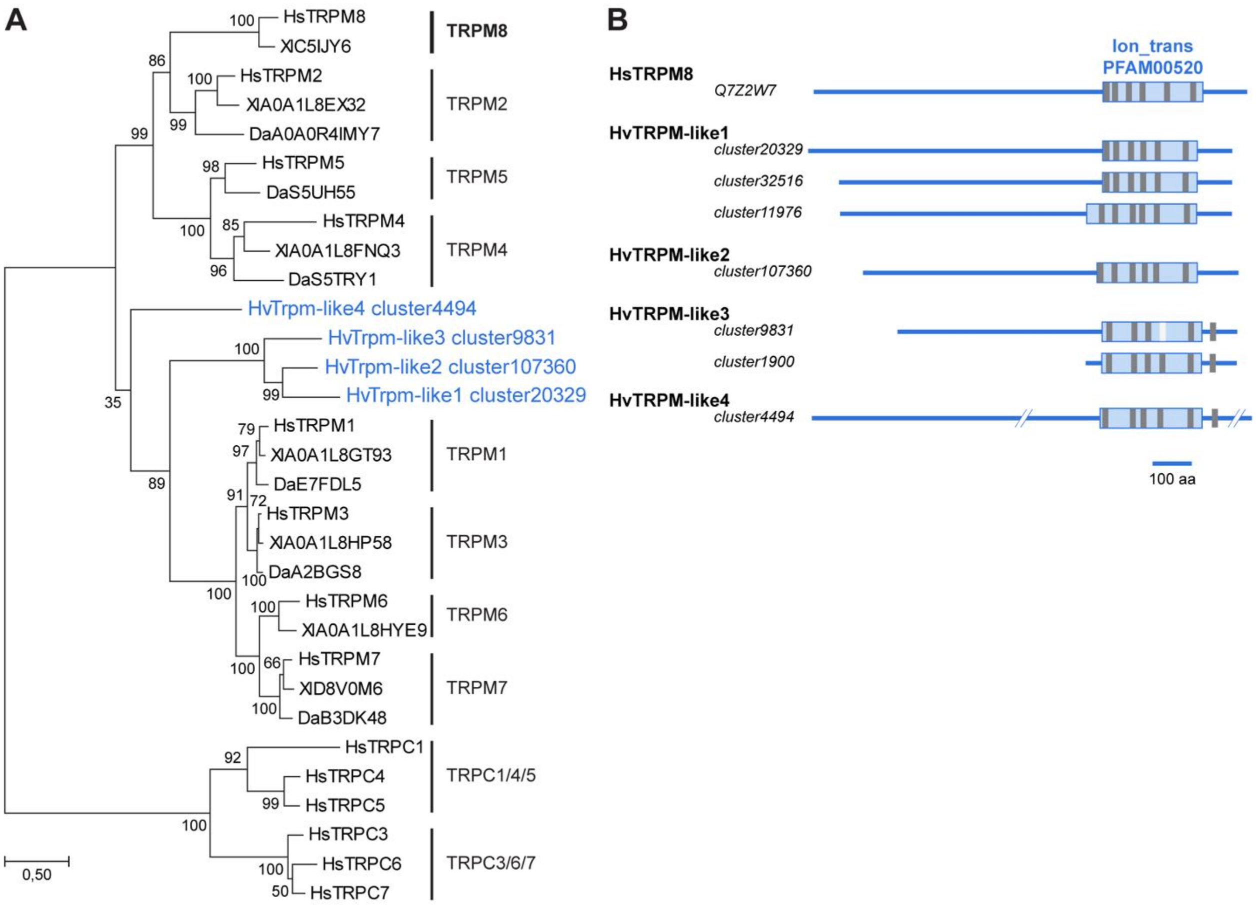


**Pacemaker-specific cation TRPM-like ion channels are highly conserved in *Hydra*. (A)** Detailed phylogenetic analysis of TRPM-like ion channels from human (*Homo sapiens*, Hs), frog (*Xenopus laevis*, Xl), zebrafish (*Danio rerio*, Da), and hydra (*Hydra vulgaris*, Hv). Human TRPC-like channels are used as an outgroup. *Hydra* has four genes coding for TRPM-like ion channels – HvTRPM-like1–4. The evolutionary history was inferred by using the Maximum Likelihood method based on the Le and Gascuel model[79]. The tree with the highest log likelihood is shown. The percentage of trees in which the associated taxa clustered together is shown next to the branches. The tree is drawn to scale, with branch lengths measured in the number of substitutions per site. Only fewer than 10% alignment gaps, missing data, and ambiguous bases were allowed at any position. There were a total of 679 positions in the final dataset. Evolutionary analyses were conducted in MEGA7[80]. (**B**) Domain structure analysis using SMART and TMHMM algorithms uncovers high conservation in structure between the human TRPM8 protein and four TRPM-like channels in *Hydra*. Multiple transcripts corresponding to the same gene were frequently identified in the reference transcriptome of *Hydra*, suggesting either existence of alternative splice-variants or poor assembly of the full-length transcripts, most likely due to the very low expression level of the genes (see Extended Data Fig. 11). Localization of specific Ion_trans (PFAM00520) domains and transmembrane domains (grey) is illustrated.

**Extended Data Fig. 10**

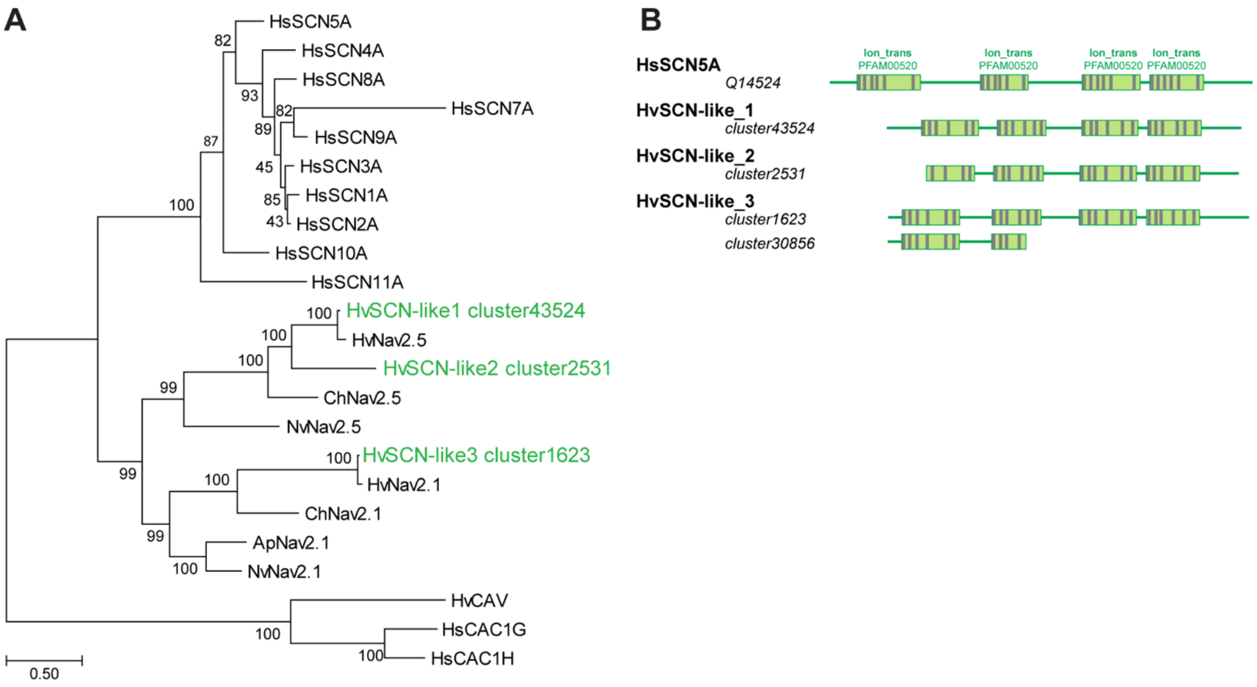


**Pacemaker-specific voltage-gated sodium SCN-like ion channels are highly conserved in *Hydra*.** (**A**) Detailed phylogenetic analysis of SCN-like ion channels from human (*Homo sapiens*, Hs), and hydra (*Hydra vulgaris*, Hv), as well as previously characterized[81] Nav channels from sea anemones (*Nematostella vectensis*, Nv, and *Aiptasia pallida*, Ap) and hydroid jellyfish (*Clytia hemisphaerica*, Ch). Human CAV and CAC channels are used as an outgroup. *Hydra* has three genes coding for SCN-like ion channels – HvSCN-like1–3. The evolutionary history was inferred by using the Maximum Likelihood method based on the Le and Gascuel model[79]. The tree with the highest log likelihood is shown. The percentage of trees in which the associated taxa clustered together is shown next to the branches. The tree is drawn to scale, with branch lengths measured in the number of substitutions per site. Only fewer than 10% alignment gaps, missing data, and ambiguous bases were allowed at any position. There were a total of 1429 positions in the final dataset. Evolutionary analyses were conducted in MEGA7[80]. (**B**) Domain structure analysis using

SMART and TMHMM algorithms uncovers high conservation in structure between the human SCN5A protein and three SCN-like channels in *Hydra*. Two transcripts corresponding to the same gene coding for HvSCN-like3 were identified in the reference transcriptome of *Hydra*, suggesting either existence of alternative splice-variants or poor assembly of the full-length transcripts, most likely due to the very low expression level of the genes (see Extended Data Fig. 11). Localization of specific Ion_trans (PFAM00520) domains and transmembrane domains (grey) is illustrated.

**Extended Data Fig. 11**

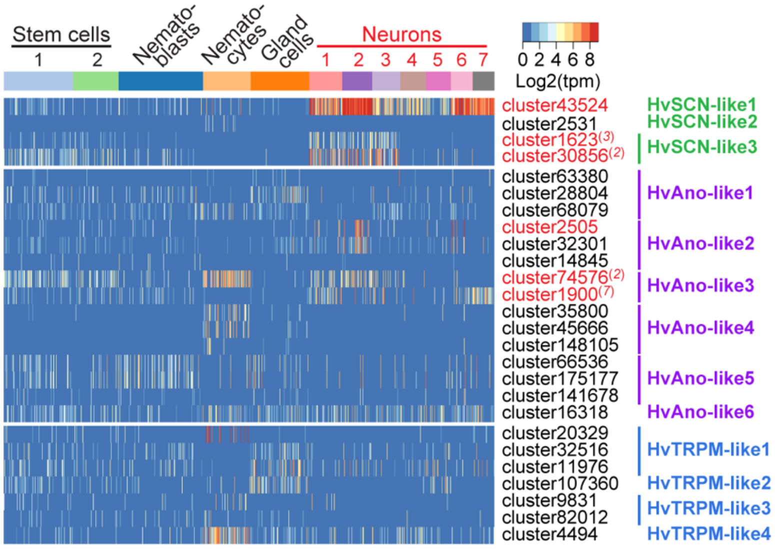


**Pacemaker-specific ion channels are expressed in *Hydra* neurons.** Heatmap illustrating expression levels of transcripts coding for putative SCN-, ANO1-, and TRPM-like ion channels in the interstitial stem cell lineage of *Hydra*. Six transcripts (marked red) are significantly upregulated in the neurons, superscript numbers indicate the nerve cell population (N1-N7) where the transcripts are significantly (padj<0.05) enriched.

**Extended Data Fig. 12**

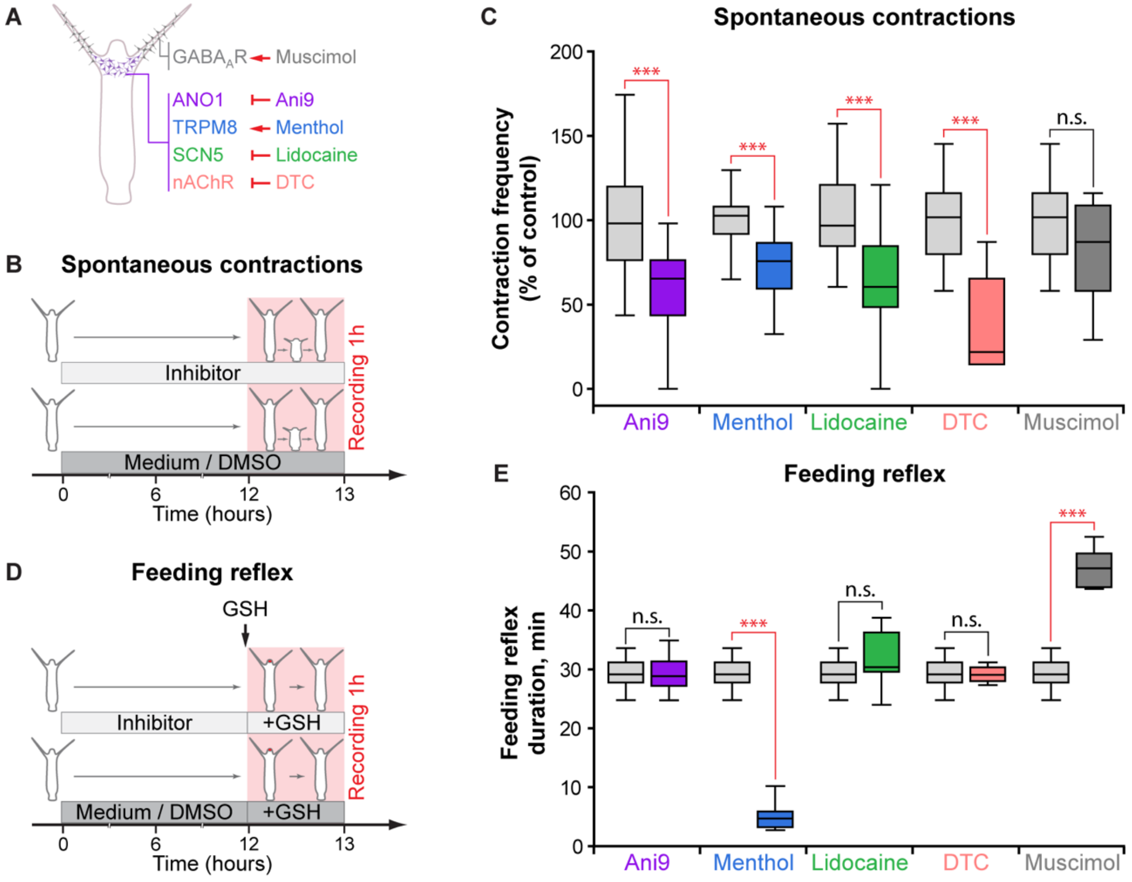


**Pharmacological substances modulate activity of ion channels and receptors and differentially affect behaviors generated by two distinct neuronal populations in *Hydra*.** (**A**) Four pharmacological agents (Ani9, Menthol, Lidocaine, and DTC) target ion channels and receptors expressed on the neuronal population N2 in the basis of tentacles. Muscimol is an agonist of $GABA_A$ receptors expressed exclusively in the neuronal population in the tentacles. (**B**) Effects of chemicals onto rhythmic spontaneous contractions of *Hydra* were assayed in the following experimental setup: polyps were incubated in the inhibitors for 12 hours prior to a 1 hour recording of contractile behavior. Polyps incubated in 0.16% DMSO-supplemented (for Ani9) or pure (all other chemicals) Hydra-medium served as control. (**C**) Contraction frequency is significantly reduced in the presence of all chemicals targeting the channels expressed on the pacemaker population N2, but not affected in the presence of muscimol, that interferes with the population N7. (**D**) Effects of pharmacological substances onto feeding reflex of *Hydra* were assayed in the following experimental setup: polyps were incubated in the inhibitors for 12 hours, the feeding reflex was induced by reduced glutathione, and the duration of feeding response (*i.e.* time between mouth opening and closure) was recorded. Polyps incubated in 0.16% DMSO-supplemented (for Ani9) or pure (all other chemicals) Hydra-medium served as control. (**E**) Three chemicals (Ani9, Lidocaine, and DTC) show no effect onto duration of the feeding reflex, while muscimol prolongs the feeding response. Complementary patterns of pharmacological effects indicate that two behaviors, the rhythmic contractions and feeding reflex, are controlled by two independent neuronal populations. Only menthol had effects onto both behaviors, reduced contraction frequency and shortened the feeding response. This may be explained by the low specificity of this chemical. In contrast to other four chemicals, that are known to be highly specific and target only narrow class of ion channels or receptors, menthol may interfere with the activity of very broad range of molecules, including ion channels, neurotransmitter receptors and enzymes[82]. Collectively, these data corroborate that the feeding reflex in *Hydra* is controlled by the GABAergic neuronal population N7, and the spontaneous contractile behavior – by the cholinergic neuronal population N2. *n*=10–49 animals (contraction frequency), *n*=11–14 animals (feeding response duration), * - $p<0.05$; ** - $p<0.005$; *** - $p<0.0005$; n.s. – $p>0.05$.

**Extended Data Fig. 13**

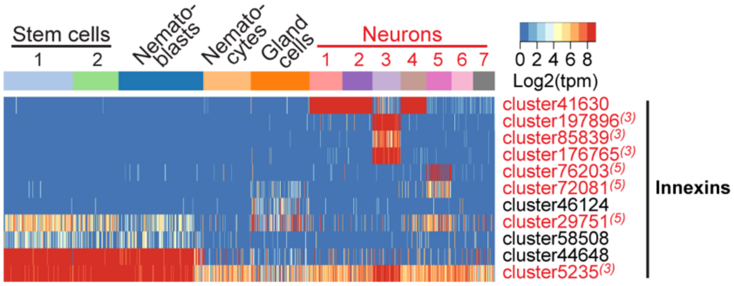


**Neurons in *Hydra* are electrically coupled by gap junctions made of innexin proteins.** Heatmap illustrating expression levels of transcripts coding for putative innexin channels in the interstitial stem cell lineage of *Hydra*. Multiple innexin genes are expressed in the neuronal populations N1–7. All cells within the population N2 that contains the pacemakers express the *innexin* transcript *cluster41630*, that is also expressed in the populations N1 and N4 localized in the foot region of a polyp (Fig. 1j). This gene codes for Innexin-2 protein that has been previously reported as essential for body contractions of *Hydra*[39].

**Extended Data Fig. 14**

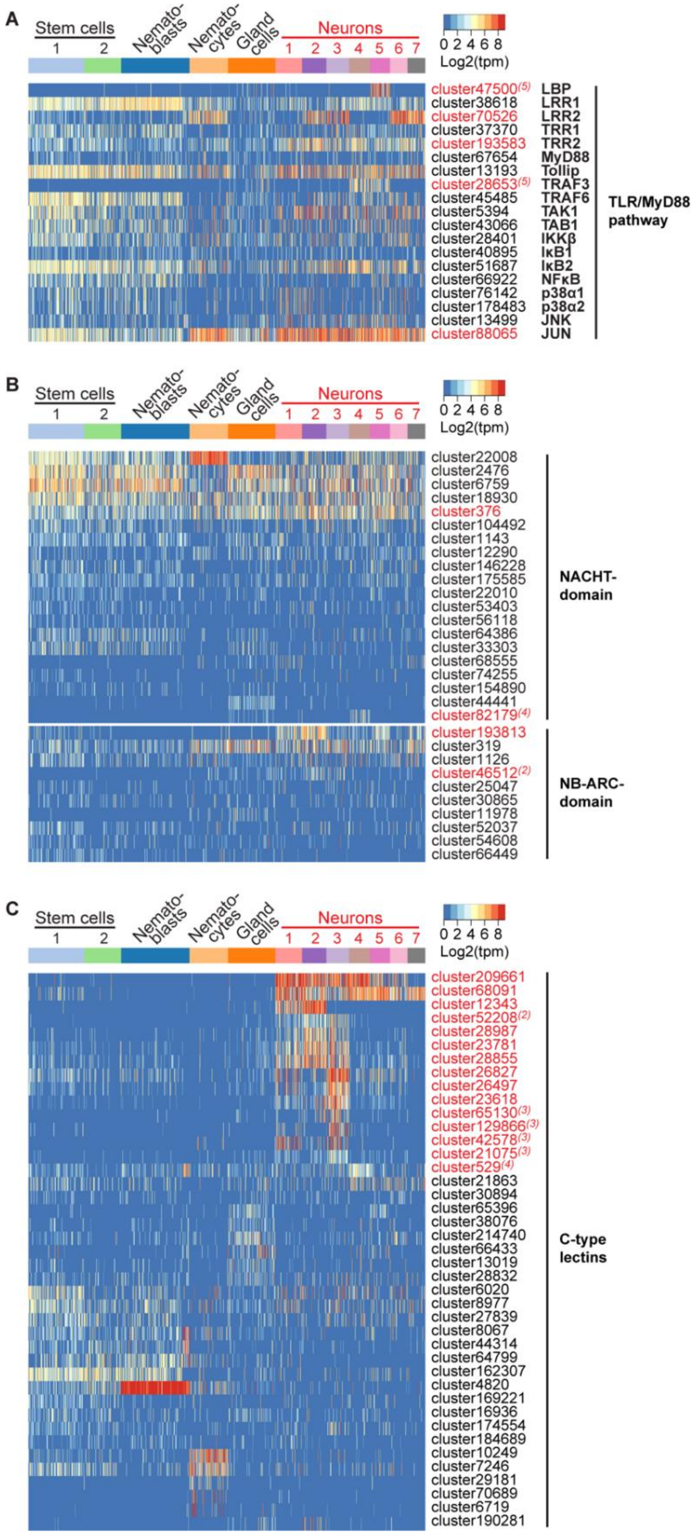

**Neurons in *Hydra* are immunocompetent cells.** (**A–C**): Heatmaps illustrating expression levels of genes coding for putative components of immune-related TLR/MyD88 pathway (on A), NACHT- and NB-ARC-domain containing NOD-like receptors (on B) and C-type lectin receptors (on C) in the interstitial stem cell lineage. Transcripts specifically upregulated in the neurons are labelled red, superscript numbers indicate the nerve cell population (N1-N7) where the transcripts are significantly ($padj<0.05$) enriched.

**Extended Data Fig. 15**

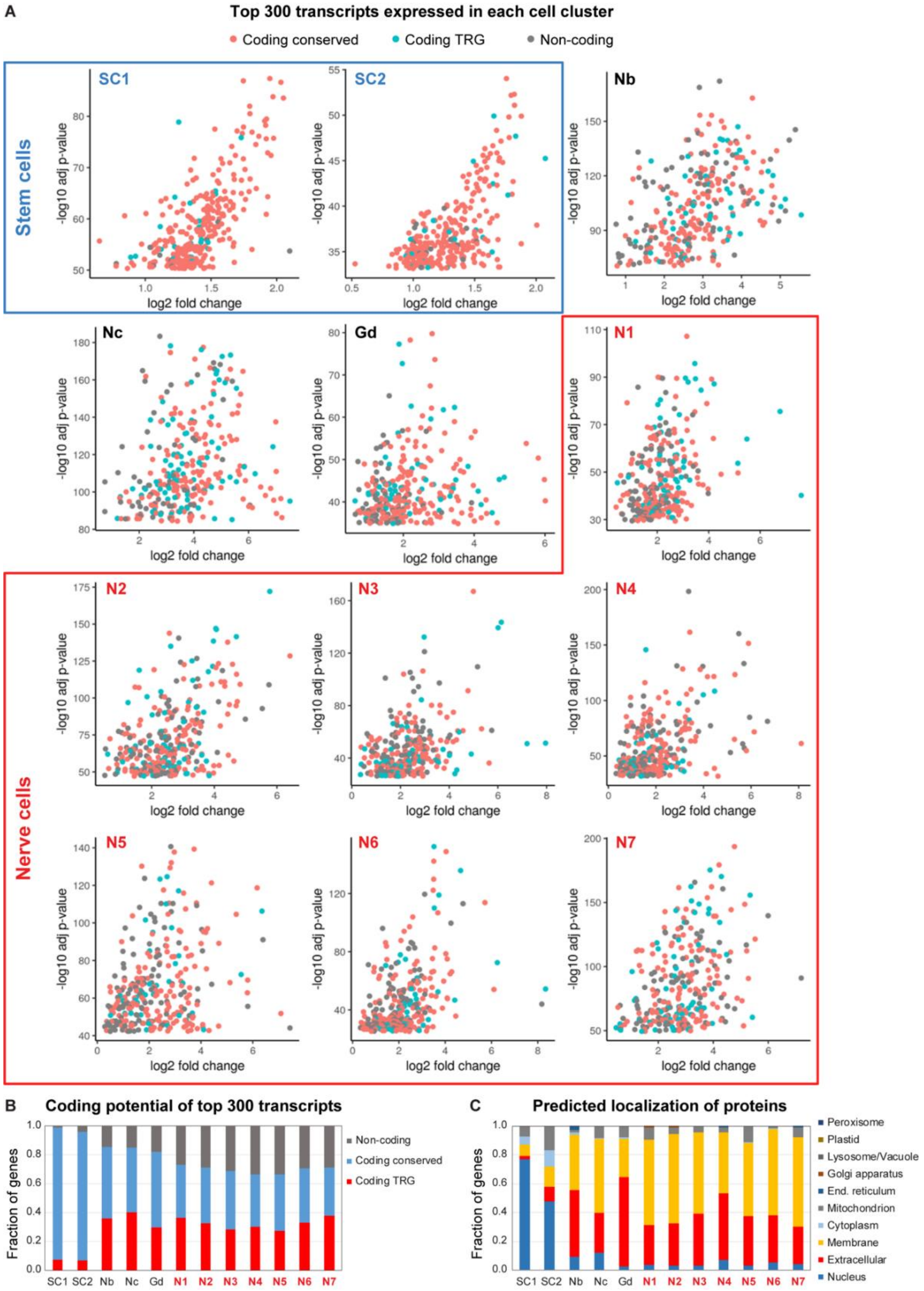

**Taxon-restricted genes (TRGs) determine the identity of each neuronal population in *Hydra* and code for predominantly secreted peptides.** (**A**) Scatter plots illustrating expression levels of top 300 genes differentially significantly up-regulated in each of 12 cell clusters. Highly-conserved protein-coding genes (torquise) dominate the signature of the stem cells. The identity of the neurons is determined by taxon-restricted protein-coding transcripts (red) and non-coding RNAs. (**B**) Relative abundance of conserved and taxon-restricted protein-coding transcripts and transcripts with no detectable open reading frame among top 300 genes differentially expressed in each of 12 clusters. (**C**) Annotation of protein-coding transcripts among top 300 genes differentially expressed in each of 12 clusters. Transcripts coding for proteins localized in the nucleus (likely, transcription factors and RNA-binding proteins; blue) dominate the signature of the interstitial stem cells. On the contrary, the contribution of transcripts encoding nuclear proteins to the identity of the neurons is minimal, consistent with our clustering based on expression of transcription factors (Extended Data Fig. 5). Instead, nearly 90% of peptides coded by transcripts specifically expressed in each neuronal population are annotated as extracellular or membrane-associated proteins (red and yellow). This suggests that the neuronal phenotype is characterized by a complex set of secreted factors (secretome), comprising mainly novel, taxon-restricted products. Some of them may mediate the communication of the nervous system with the microbiota.

**Extended Data Fig. 16**

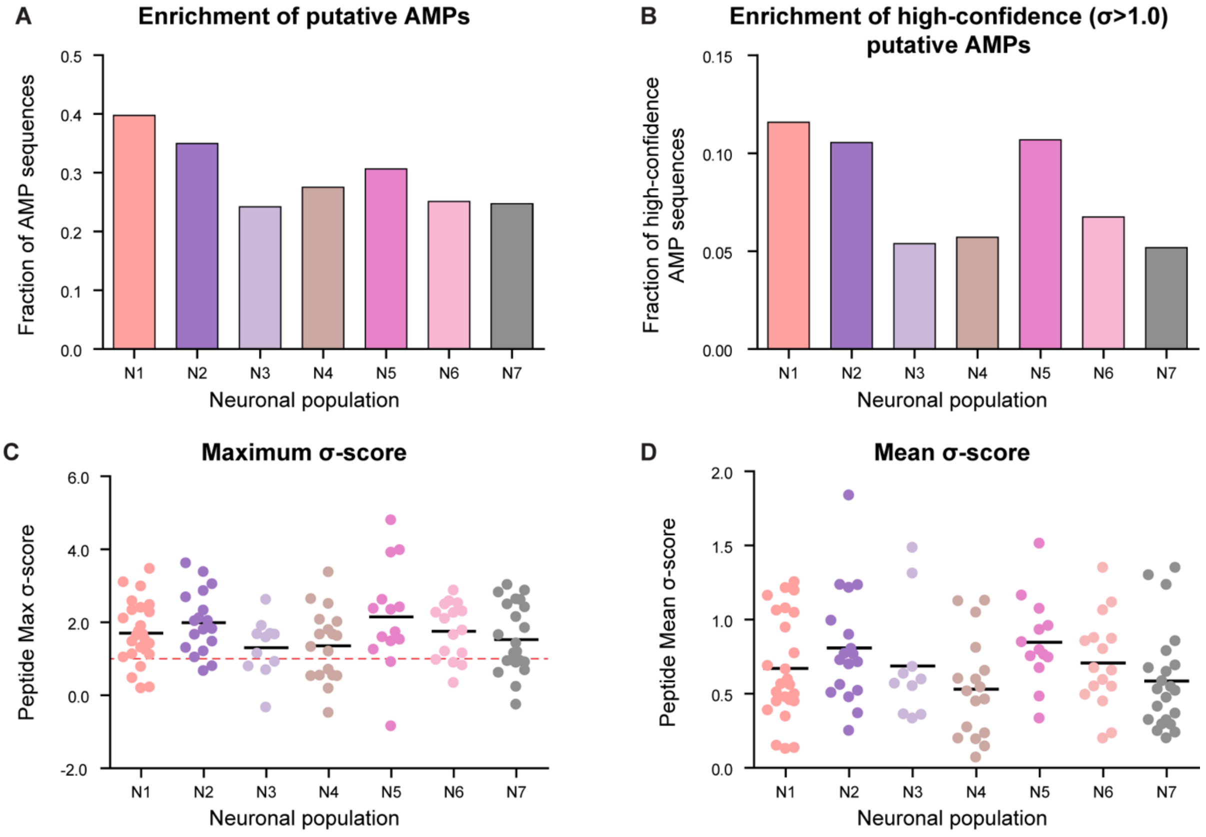


**Secreted peptides encoded in neuron-specific TRGs contain putative antimicrobial peptides.** **(A)** A substantial fraction of secreted peptides encoded by neuron-specific TRGs contains small peptides that may function as antimicrobial peptides, as revealed by a moving window (20 aa) small-peptide scan. (**B**) Neuronal populations N1, N2 and N5 are particularly enriched in putative AMPs with high confidence (σ>1.0) membrane destabilizing activity. (**C**) Distribution of maximum σ-score values for each of secreted peptides encoded by neuron-specific TRGs in seven neuronal subpopulations. Raw values are provided in Extended Data 3. Majority of the TRG-encoded peptides contain at least one subsequence of 20 aa with a positive σ-score. The neuronal population N2 that contains the pacemakers is particularly rich in secreted peptides with high-confidence (σ>1.0, red dotted line) antimicrobial activity. (**D**) Distribution of mean σ-score values for each of secreted peptides encoded by neuron-specific TRGs in seven neuronal subpopulations illustrates a high likelihood of containing an AMP for the peptides. Raw values are provided in Extended Data 3.

**Extended Data Fig. 17**

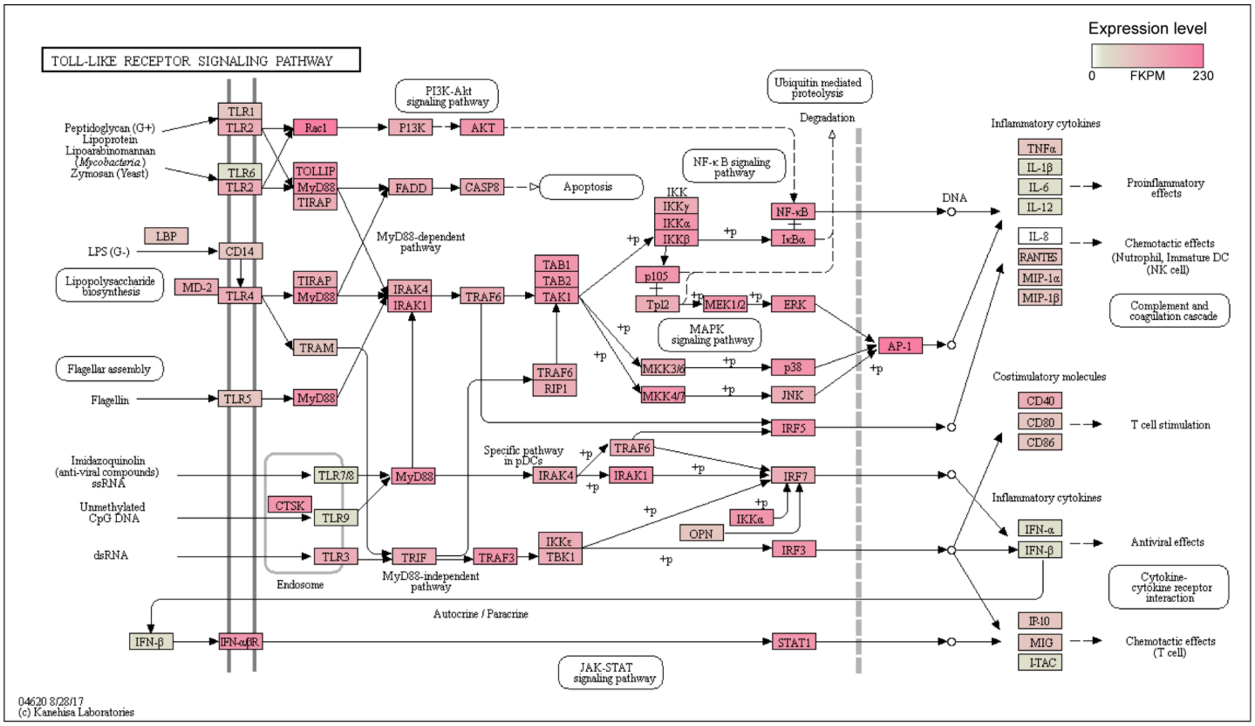


**Murine pacemaker cells, the interstitial cells of Cajal (ICC), express immune-related TLR/MyD88 pathway components.** KEGG-pathway mapping. Genes highly abundant in the transcriptome of ICC cells[15] are marked red, poorly expressed genes in gray, transcripts missing in the dataset are labeled white. Original data are presented in Extended Data 4.

**Extended Data Fig. 18**

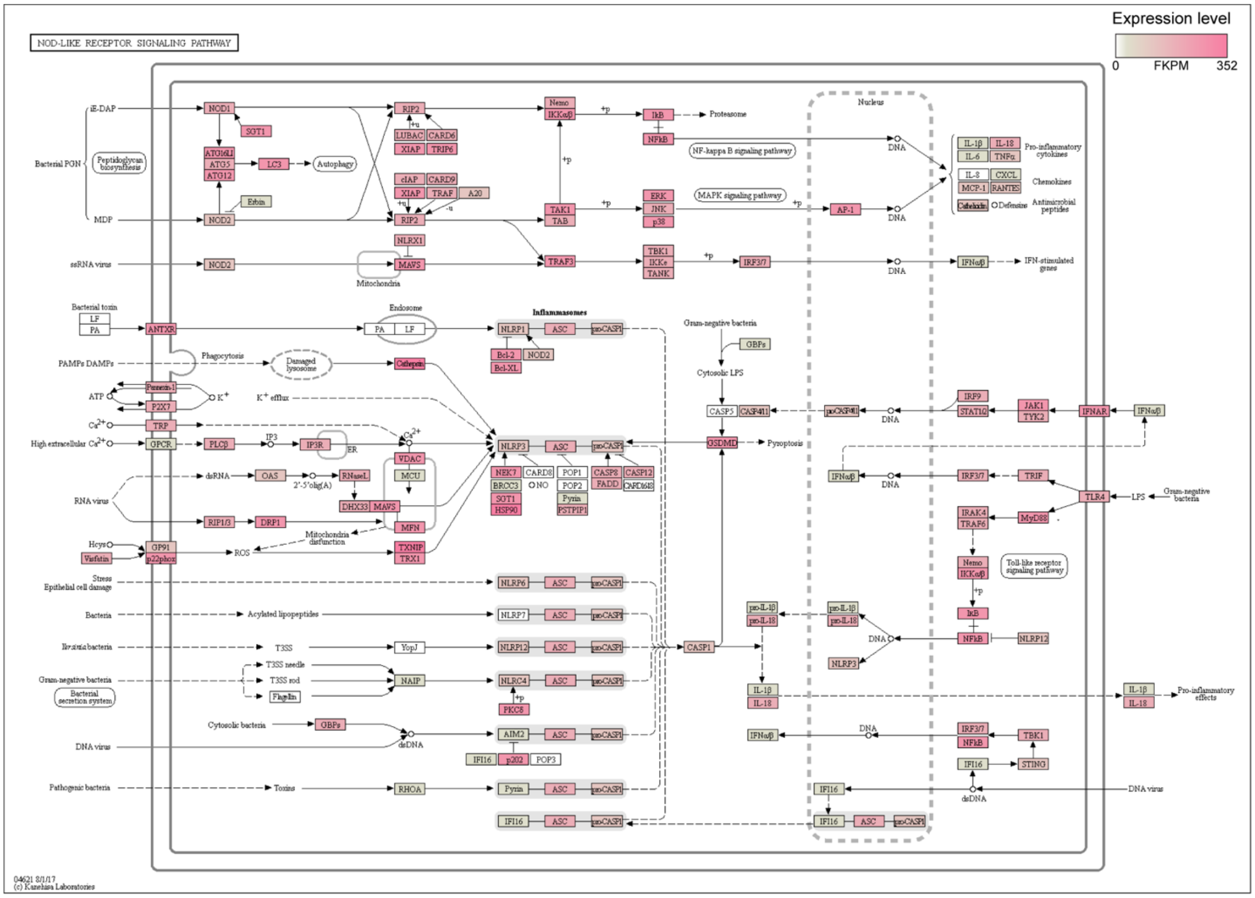


**Murine pacemaker cells, the interstitial cells of Cajal (ICC), express immune-related NOD-like receptor pathways.** KEGG-pathway mapping. Genes highly abundant in the transcriptome of ICC cells[15] are marked red, poorly expressed genes in gray, transcripts missing in the dataset are labeled white. Original data are presented in Extended Data 4.

**Extended Data Fig. 19**

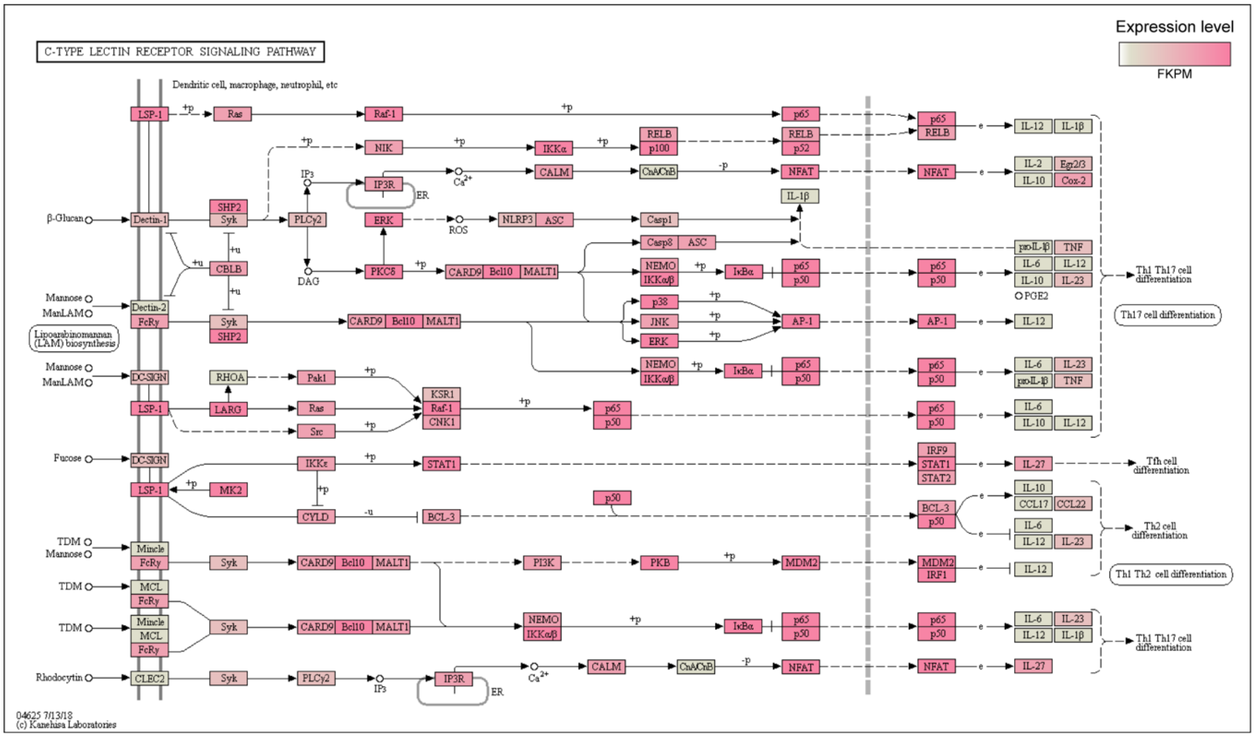


**Murine pacemaker cells, the interstitial cells of Cajal (ICC), express immune-related C-type lectin pathway components.** KEGG-pathway mapping. Genes highly abundant in the transcriptome of ICC cells[15] are marked red, poorly expressed genes in gray, transcripts missing in the dataset are labeled white. Original data are presented in Extended Data 4.

**Supplementary Table 1**

Oligonucleotide primers used to amplify gene fragments in qRT-PCR.

| Target gene | **Forward primer 5’ -> 3’** | **Reverse primer 5’ -> 3’** |
|---|---|---|
| *actin* | gaatcagctggtatccatgaaac | aacattgtcgtaccacctgatag |
| *elongation factor 1* | gcagtactggtgagtttgaag | cttcgctgtatggtggttcag |
| *cluster43524* | *tggtggtgatgcgctgag* | *tcgatcgactttgtttgaccc* |
| *cluster63380* | *tgcttgcacgggttggac* | *gtagtttcaggtagacacagg* |
| *cluster28804* | *ctgagcagcacgatcgtg* | *aggcgacattgctggctc* |
| *cluster2505* | *gctccattagcgctacgtc* | *aagtagtcaggaggactcatag* |
| *cluster32301* | *gaccaagggattcaagaagag* | *ccatctcgaaagtaagtccc* |
| *cluster74576* | *tcaaactgatcgctcaagaattg* | *ccatcatgaacaggataacaatc* |
| *cluster11976* | *aaacgaagagacaagcttagttg* | *tctgttggtgtctcttgtgg* |

**Extended Data 1. (separate file)**

Dataset of 364 transcripts coding for putative transcription factors.

**Extended Data 2. (separate file)**

Dataset of top 300 transcripts expressed in each of 12 populations within the interstitial cell lineage. Cluster numbers, best hit annotation using UniProt and RefSeq databases, predicted peptide length and domains, signal peptide and cellular localization.

**Extended Data 3. (separate file)**

Dataset of putative secreted antimicrobial peptides encoded by neuron-specific TRGs. Predicted membrane destabilizing activity estimated using machine learning tool[49] with a moving window of 20 amino acids.

**Extended Data 4. (separate file)**

Expression of genes coding for components of immune-related TLR/MyD88 pathways, NOD-like and C-type lectin receptors in the murine intestinal pacemaker cells, interstitial cells of Cajal (ICC).

**Extended Data 5. (separate file)**

Dataset of 25 transcripts comprising the proliferation signature used for annotation of cell clusters.

**Extended Data 6. (separate file)**

Dataset of 24 transcripts of cell-type specific marker genes used for annotation of cell clusters.

**Extended Data 7. (separate file)**

Dataset of 112 transcripts coding for putative neurotransmitter receptors.

**Extended Data 8. (separate file)**

Dataset of 431 transcripts coding for putative ion channels.
